# Identification of novel disease relevant genetic modifiers affecting the SHH pathway in the developing brain

**DOI:** 10.1101/2020.11.03.366302

**Authors:** Nora Mecklenburg, Izabela Kowalczyk, Franziska Witte, Jessica Görne, Alena Laier, Hannes Gonschior, Martin Lehmann, Matthias Richter, Anje Sporbert, Bettina Purfürst, Norbert Hübner, Annette Hammes

## Abstract

Pathogenic gene variants in humans affecting the sonic hedgehog (SHH) pathway lead to severe brain malformations with variable penetrance due to unknown genetic modifiers. To identify such modifiers, we established novel congenic mouse models. LRP2 deficient C57BL/6N mice suffer from heart outflow tract defects and holoprosencephaly caused by impaired SHH activity. These defects are fully rescued on FVB/N background indicating a strong influence of modifier genes. Applying comparative transcriptomics, we identified *Pttg1* and *Ulk4* as candidate modifiers upregulated in the rescue strain. Functional analyses showed that ULK4 and PTTG1, both microtubule-associated proteins, are new positive regulators of SHH signaling, rendering the pathway more resilient to disturbances. In addition, we characterized PTTG1 as a novel primary cilia component in the neuroepithelium. The identification of genes, that powerfully modulate the penetrance of genetic disturbances affecting the brain and heart, is likely relevant to understand variability in human congenital disorders.

## INTRODUCTION

Holoprosencephaly (HPE) is the most common structural defect of human forebrain development resulting from impaired separation of the two cephalic hemispheres and typically accompanied by craniofacial malformations (Geng and Oliver, 2009; Hong and Krauss, 2018; Krauss, 2007; Ming and Muenke, 2002; Muenke and Beachy, 2000). HPE is a complex condition caused by genetic predisposition and exposure to environmental toxins (Hong et al., 2020; Krauss and Hong, 2016; Tekendo-Ngongang et al., 1993). The genetic contribution can be either monogenic or polygenic (Krauss and Hong, 2016; Roessler and Muenke, 2010). Single mutations causing HPE have been identified in mice and humans (Hayhurst and McConnell, 2003; Roessler and Muenke, 2010; Roessler et al., 1996). Mutations in the human sonic hedgehog (*SHH*) gene and its downstream effector genes account for at least 5% of autosomal dominant nonsyndromic HPE cases (Wallis and Muenke, 2000, 1999; Wallis et al., 1999). Mice deficient for SHH develop forebrain defects resembling those of human HPE cases (Chiang et al., 1996). Another gene more recently implicated in the etiology of HPE in mice and humans is *LRP2* (low-density lipoprotein receptor-related protein 2) (Kim et al., 2019; Rosenfeld et al., 2010; Spoelgen et al., 2005; Willnow et al., 1996). Loss of LRP2 activity in *Lrp2*^−/−^ mouse embryos causes HPE, associated with severe craniofacial defects (Spoelgen et al., 2005; Willnow et al., 1996). LRP2 is also a component of the SHH signaling machinery in the primary cilium (Christ et al., 2012), as LRP2 deficiency leads to an impairment in the response of the ventral forebrain neuroepithelium to SHH.

Humans with autosomal recessive *LRP2* gene defects suffer from Donnai-Barrow syndrome with clinical characteristics resembling those in the mouse model, including craniofacial anomalies, forebrain defects (Kantarci et al., 2007; Ozdemir et al., 2019), microforms of HPE (Rosenfeld et al., 2010), and heart anomalies (Pober et al., 2009). A recent publication describes severe forms of HPE in families presenting oligogenic events involving *LRP2* (Kim et al., 2019). Inter- and intrafamilial phenotypic variability is observed in cases of the Donnai-Barrow syndrome (Khalifa et al., 2015; Longoni et al., 1993; Pober et al., 2009) and is a characteristic feature of HPE (Dubourg et al., 2018; Solomon et al., 2010). Even within pedigrees, HPE phenotypes amongst relatives, carrying the same *SHH* mutation, can range from alobar HPE, facial abnormalities typical of HPE, to asymptomatic appearance of the carrier. Intrafamilial variability of HPE phenotypes could be due to environmental or genetic factors. (Heussler et al., 2002; Hong and Krauss, 2018; Ming and Muenke, 2002; Muenke and Beachy, 2000; Muenke and Cohen, 2000; Roessler et al., 1996).

Studies on mouse models for HPE suggest that loss of function mutations in genes relevant for forebrain development result in brain and craniofacial anomalies that recapitulate HPE phenotypes in patients carrying mutations in the same genes (Geng and Oliver, 2009; Hayhurst and McConnell, 2003; Heyne et al., 2016; Hong and Krauss, 2018). Intriguingly, the variability of the HPE expression is also reflected in gene targeted mouse models since the penetrance of the phenotype often depends on the mouse strain that is analyzed (Anderson et al., 2002; Cole and Krauss, 2003; Geng and Oliver, 2009; Geng et al., 2008; Hong and Krauss, 2018; Petryk et al., 2004; Schachter and Krauss, 2008). Thus, the mouse is an ideal model, not only to investigate the complex etiology of human HPE but also to identify disease relevant genetic modifiers.

## RESULTS

### Genetic background determines penetrance of brain, craniofacial and heart defects in *Lrp2*^−/−^ mutant mice

*Lrp2*^−/−^ mice on a C57BL/6N background at embryonic stage (E) 18.5 displayed severe HPE associated skull and facial dysgenesis (Figure 1A: d - f). In particular, they showed a shortened snout (Figure 1A: d), cleft lip (Figure 1A: e, arrow) and incompletely developed or even bilaterally or unilaterally absent eyes (Figure 1A: d and e). Further characteristic features seen in these mutants were a shortened skull (Figure 1A: f) and an open anterior suture (Figure 1A: f, arrowhead). In contrast, *Lrp2*^−/−^ congenic mice with a FVB/N background showed 100% penetrant rescue of all the above described craniofacial defects (Figure 1A: j – l). HPE in *Lrp2*^−/−^ C57BL/6N embryos was fully penetrant and characterized by a fusion of the cortical hemispheres (Figure 1B: d, arrowhead) with a single lateral ventricle (Figure 1B: f, arrowhead) and absent olfactory bulbs (Figure 1B: d and e, asterisk). In contrast, HPE was fully rescued in all *Lrp2*^−/−^ FVB/N mutants, which presented with correctly separated cortical hemispheres, olfactory bulbs and clearly defined lateral ventricles (Figure 1B: j – l). Corpus callosum and the anterior commissure were also present in the LRP2 deficient mice on a FVB/N background (Figure 1B: l).

**Figure 1:**
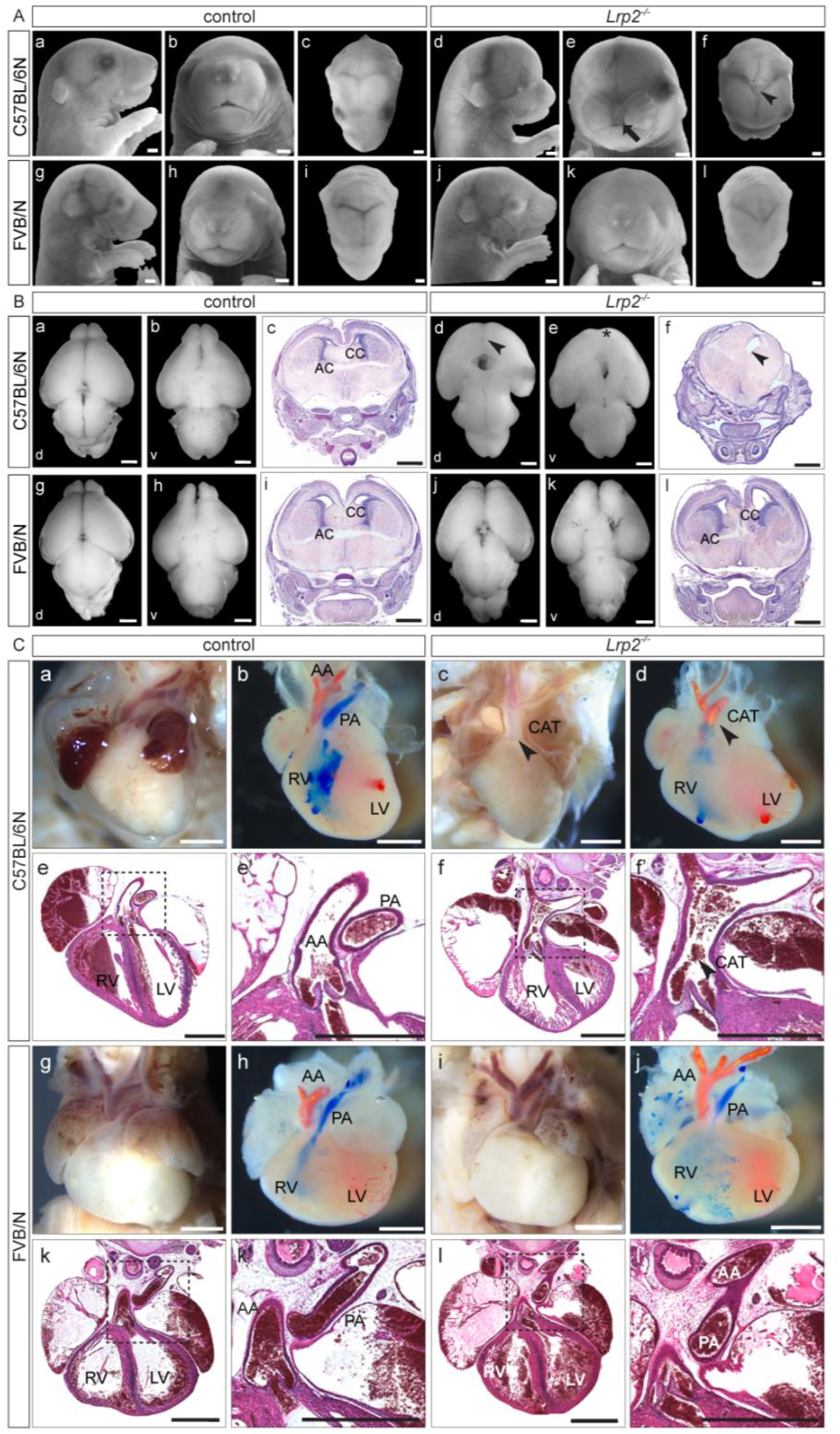
Rescue of craniofacial malformations, HPE and heart defects in *Lrp2*^−/−^ FVB/N mice. (**A**) *Lrp2*^−/−^ C57BL/6N E18.5 embryos (n = 30) with craniofacial defects: shortened snout (**d**), underdeveloped or missing eyes (**e**), cleft lip (**e**, arrow) and hypertelorism (**e**). Dorsal view on the head revealed shortened and widened skull with impaired suture formation and an open fontanelle (**f**, arrowhead). *Lrp2*^+/+^ and *Lrp2*^+/−^ littermates (**a**-**c**) served as controls (n = 66). *Lrp2*^−/−^ FVB/N mutants (n = 28) displayed normal skull formation (**j**), normal eyes and normal midface structures (**k**) comparable to controls (n = 94) (**g**-**i**). Note, missing pigmentation of retina due to albinism in FVB/N. Scale bars: 1 mm (**B**) Brains from E18.5 control and *Lrp2*^−/−^ embryos in dorsal and ventral view and NISSL stained coronal sections. *Lrp2*^−/−^ C57BL/6N mice with HPE (**d**-**f**). *Lrp2*^−/−^ C57BL/6N brain with alobar HPE (**d**, arrowhead) (*Lrp2*^−/−^ n = 4; controls n = 6). Olfactory bulbs were absent in mutants (**e**, asterisk). All sectioned *Lrp2*^−/−^ C57BL/6N brains (n = 13) showed alobar HPE with a single lateral ventricle (**f**, arrowhead) compared to controls (**c**), (n = 6). Missing corpus callosum (CC) and anterior commissure (AC) in *Lrp2*^−/−^ C57BL/6N mice (**f**). *Lrp2*^−/−^ FVB/N embryos showed normal forebrain separation (**j**-**l**); (*Lrp2*^−/−^ n = 5; controls n = 3). Olfactory bulbs were present in *Lrp2*^−/−^ FVB/N brains (**k**). All *Lrp2*^−/−^ FVB/N sectioned brains (n = 6) displayed normally separated ventricles, normal CC and AC (**l**) comparable to controls (**i**), (n = 4). Scale bars: 1 mm. d: dorsal, v: ventral. (**C**) Analysis of heart outflow tract phenotype in *Lrp2*^−/−^ mutants and controls. C57BL/6N (**a** and **b**) and FVB/N (**g** and **h**) control (refers to *Lrp2*^+/+^ and *Lrp2*^+/−^) hearts at E18.5 by external inspection and pigment injection. In controls (**b** and **h**) injection of Batson’s red pigment into the left ventricle after cutting the ductus arteriosus indicated the ascending aorta in red (AA). Injection of blue pigment into the right ventricle labeled the pulmonary artery in blue (PA). Frontal plane H&E stained paraffin sections through hearts of control embryos (**e, e’** and **k**, **k’**) depicted normally separated ascending aorta (AA) and pulmonary artery (PA). *Lrp2*^−/−^ mutants (15/16) on C57BL/6N background suffered from common arterial trunk (CAT), (**c**, arrowhead). 1/16 mutants suffered from double outlet right ventricle (DORV, see **Supplementary Table 1**). Injection of the red pigment into the mutant’s left ventricle followed by injection of blue ink into the right ventricle showed both pigments in the CAT and indicated an additional ventricular septum defect (**d,** arrowhead). CAT in *Lrp2*^−/−^ C57BL/6N embryos was also shown on H&E stained frontal heart sections (**f**, **f’**) No CAT was detected in *Lrp2*^−/−^ mutants on FVB/N background (**i, j, l, l’**). 85% of *Lrp2*^−/−^ FVB/N embryos displayed a normal heart morphology with a proper connection of the ascending aorta to the left ventricle and of the pulmonary artery to the right ventricle. 2 out of 13 *Lpr2*^−/−^ FVB/N mice showed DORV (**Supplementary Table 1**). Scale bars: 1 mm. LV: left ventricle; RV: right ventricle.

LRP2 deficient mice also suffer from cardiovascular anomalies including a common arterial trunk (CAT) as one of the most penetrant phenotypes on a C57BL/6N background (Baardman et al., 2016; Christ et al., 2020; Li et al., 2015). Interestingly, there is comorbidity of HPE and congenital heart disease in patients with variants of known HPE-related genes (Tekendo-Ngongang et al., 2020). *Lrp2*^−/−^ embryos with CAT, also known as persistent truncus arteriosus, displayed a single common coronary outflow artery vessel instead of pulmonary artery and ascending aorta. Less frequently *Lrp2*^−/−^ embryos displayed a double outlet right ventricle (DORV), in which the pulmonary artery and ascending aorta both connect to the right ventricle. We found that 15 out of 16 *Lrp2*^−/−^ mutants on a C57BL/6N background displayed CAT above the right ventricle (Figure 1C: c, d, f, f’ and Supplementary Table 1), and one remaining mutant embryo displayed DORV (Supplementary Table 1). The CAT phenotype was also visualized by pigment injection, (Figure 1C: d, arrowhead) indicating CAT and a ventricular septum defect. We never observed CAT in *Lrp2*^−/−^ FVB/N mutants; 85% of *Lrp2*^−/−^ FVB/N embryos showed no apparent heart anomalies and a normal ventricular septum compared to control littermates (Figure 1C: i, j, l, l’ and Supplementary Table 1). Two out of 13 *Lrp2*^−/−^ FVB/N mice showed a double outlet right ventricle (DORV), (Supplementary Table 1), which has been described in mutants on C57BL/6N strain with a similar frequency (Baardman et al., 2016). We conclude that the FVB/N background fully rescues the CAT phenotype, whereas the etiology of DORV is not influenced by the strain background, suggesting the likelihood of divergent pathogenic mechanisms for CAT and DORV in *Lrp2*^−/−^ mutants.

### Rescue of SHH activity in neuroepithelial stem cells

In the neuroepithelium, LRP2 is located at the base of the primary cilium, where it teams up with Patched1 to mediate endocytic uptake of SHH and most importantly recycling of the morphogen (Christ et al., 2012). Consistent with our previous findings, *Lrp2*^−/−^ C57BL/6N embryos showed normal SHH pattern in the prechordal plate, but little to no SHH in the overlying neuroepithelial stem cells compared to control embryos during the critical stages of forebrain specification at the 8 somite stage (Figure 2B). SHH protein was detected later in the neuroepithelium at the 11 somite stage, which is too late for proper ventral midline specification (Figure 2B). Intriguingly, all *Lrp2*^−/−^ FVB/N mutants showed completely normal SHH localization throughout all critical developmental stages (Figure 2B). To exclude later onset defects, we analyzed later embryonic stages. A fully penetrant hallmark of the *Lrp2*^−/−^ C57BL/6N embryos at E10.5 is the lack of *Shh* RNA and protein in the telencephalon (Christ et al., 2012; Spoelgen et al., 2005) (Figures 2C, arrowheads and S2A: b) and in the ventral midline of the diencephalon (Figure S2B: b). In contrast, *Lrp2*^−/−^ FVB/N mutants showed normal *Shh* RNA expression and SHH protein localization in the ventral telencephalon and diencephalon (Figures 2C, arrowhead, S2A: d, and S2B: d).

**Figure 2:**
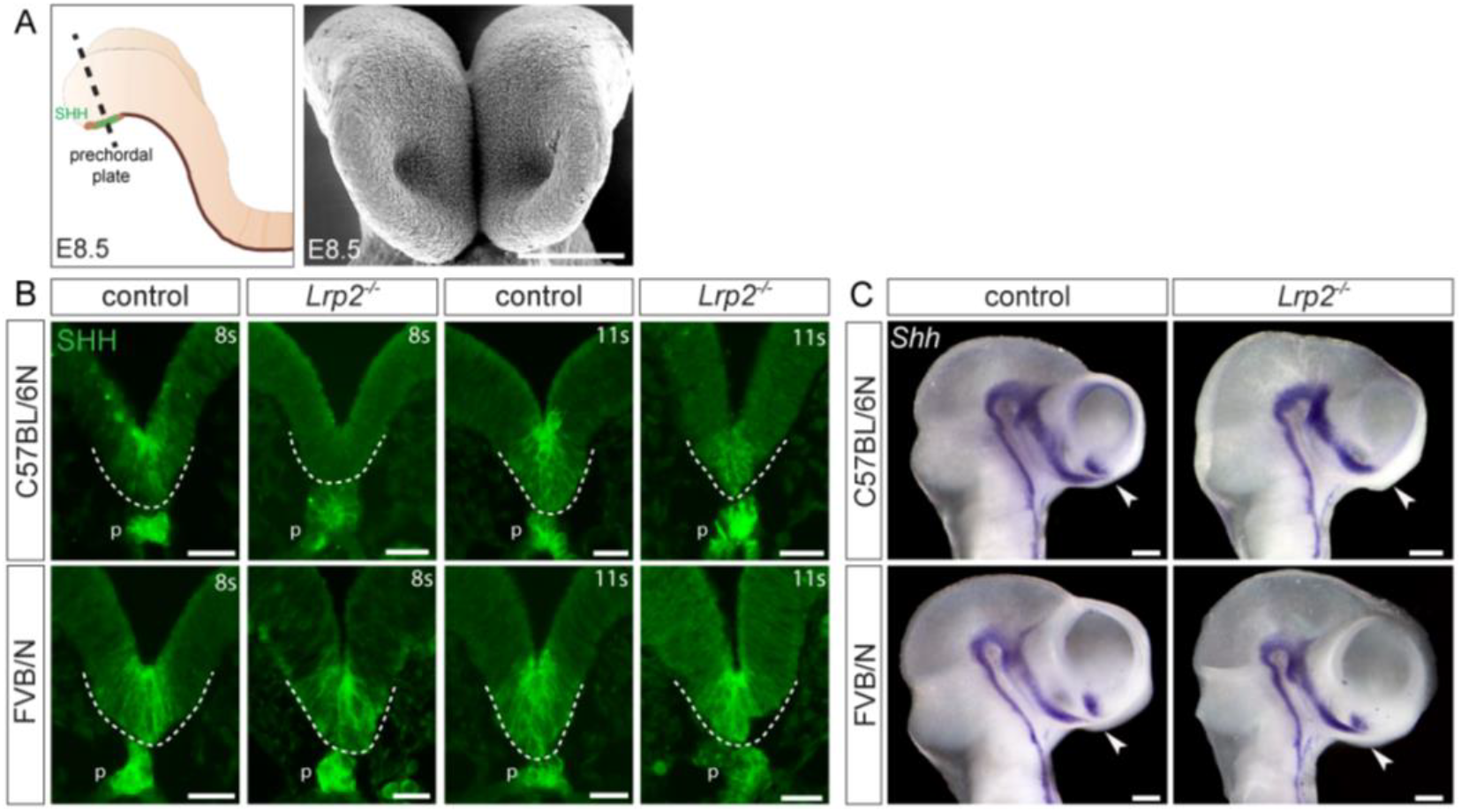
Loss of SHH is rescued in the ventral forebrain of *Lrp2*^−/−^ FVB/N mutants. (**A**) Schematic of E8.5 mouse embryo indicating the prechordal plate, underlying the neuroepithelium. Dotted line indicates section planes shown in (**B**). Scanning electron microscopy image represents the front view on the developing neural folds. Scale bar: 100 μm. (**B**) Immunohistology for SHH on coronal sections of anterior neural folds in control (*Lrp2^+/+^* and *Lrp2*^+/−^) and *Lrp2*^−/−^ embryos on C57BL/6N and FVB/N background. In 8 somite stage (8 s) control embryos (n = 7), SHH was detected in the prechordal plate and in the neuroepithelium (above the dotted line). *Lrp2*^−/−^ C57BL/6N embryos at 8 s (n = 4), showed SHH in the prechordal plate, but little to none SHH in the neuroepithelium. In 11 somite stage embryos SHH appeared in the neuroepithelium of mutant embryos (n = 3; n = 4 for controls). *Lrp2*^−/−^ FVB/N embryos (n = 8 at 8 - 9 s, n = 2 at 11 s) displayed normal SHH pattern in the neuroepithelium throughout development, comparable to somite matched control samples (n = 10 at 8 - 9 s, n = 8 at 11 s). Scale bars: 25 μm. p: prechordal plate, s: somites. (**C**) Whole mount ISH hybridization for *Shh* on E10.5 embryonic heads. Controls (n = 5) showed typical *Shh* expression pattern with a prominent signal in the ventral telencephalon (arrowhead). *Lrp2*^−/−^ C57BL/6N somite matched embryos (n = 9) had no telencephalic *Shh* signal (arrowhead). *Lrp2*^−/−^ FVB/N embryos (n = 8) displayed the same expression pattern as controls (n = 4). Scale bars: 250 μm. See also Figure S2.

Normal *Nkx2.1* expression pattern comparable to controls (Figure S2C: a, b, e, f) confirmed proper activation of SHH downstream targets and intact ventral forebrain patterning in *Lrp2*^−/−^ FVB/N mice (Figure S2C: g, h) in contrast to mutants on a C57BL/6N background, which lacked *Nkx2.1* expression in the ventral forebrain (Figure S2C: c, d).

Thus, defects in the SHH pathway caused by loss of LRP2 were fully rescued on a FVB/N background.

Consequently, the severe telencephalic vesicle separation defects in *Lrp2*^−/−^ C57BL/6N mutants (Figure S2D: c, d) did not manifest in mutants on a FVB/N background displaying properly separated forebrain hemispheres at E12.5 (Figure S2D: g, h).

### RNA sequencing reveals a dominant effect of FVB/N specific rescue modifier genes

We next analyzed the transcriptome of *Lrp2^+/+^* and *Lrp2*^−/−^ embryonic heads from C57BL/6N and FVB/N backgrounds (Figure 3A). To test a possible dominant effect of FVB/N specific modifier genes, we also collected *Lrp2^+/+^* and *Lrp2*^−/−^ first generation hybrids from *Lrp2*^+/−^ C57BL/6N x *Lrp2*^+/−^ FVB/N parents, hereon referred to as F1 samples, for RNA sequencing (Figure 3A). Performing DESeq2 differential expression analyses on all transcriptomes (Figure 3B heatmap and Supplementary Table 2), we detected 2170 differentially expressed genes (DEGs) between *Lrp2*^−/−^ and *Lrp2^+/+^* embryos on a C57BL/6N background and only 367 DEGs for the FVB/N background and a similarly low number (241 DEGs) for the F1 background (Figure 3C). Expression levels of HPE- and SHH pathway-related transcription factors like *Vax1*, *Six6* and *Six3* were significantly decreased in *Lrp2*^−/−^ C57BL/6N mutants compared to controls, but were unchanged in *Lrp2*^−/−^ FVB/N mutants compared to FVB/N controls (Figure 3D). Interestingly, also the *Lrp2*^−/−^ F1 mutants showed normal expression levels for *Vax1*, *Six6* and *Six3*, comparable to the *Lrp2^+/+^* controls (Figure 3D). Detailed phenotypic analyses of the *Lrp2*^−/−^ F1 embryos indeed revealed a full rescue of brain and craniofacial defects in these F1 mutants (Figures 3E and S3A: e – h). They neither suffered from HPE (Figures 3E and S3A: h) nor cleft lip (Figure S3A: f). Analyzing the heart, we never observed a common outflow tract (0/27) and 56% of the *Lrp2*^−/−^ mutant F1 embryos had normal hearts and outflow tracts (Figure S3B: d - f and Supplementary Table 1). However, in 44% of mutants we detected a double outlet right ventricle (DORV), where pulmonary artery and ascending aorta both connect to the right ventricle (Figure S3B: g - i). Most importantly, as observed for the *Lrp2*^−/−^ FVB/N embryos, all *Lrp2*^−/−^ F1 embryos (8/8) showed normal *Shh* expression in the developing forebrain similar to control embryos (Figure 3F, arrowheads).

**Figure 3:**
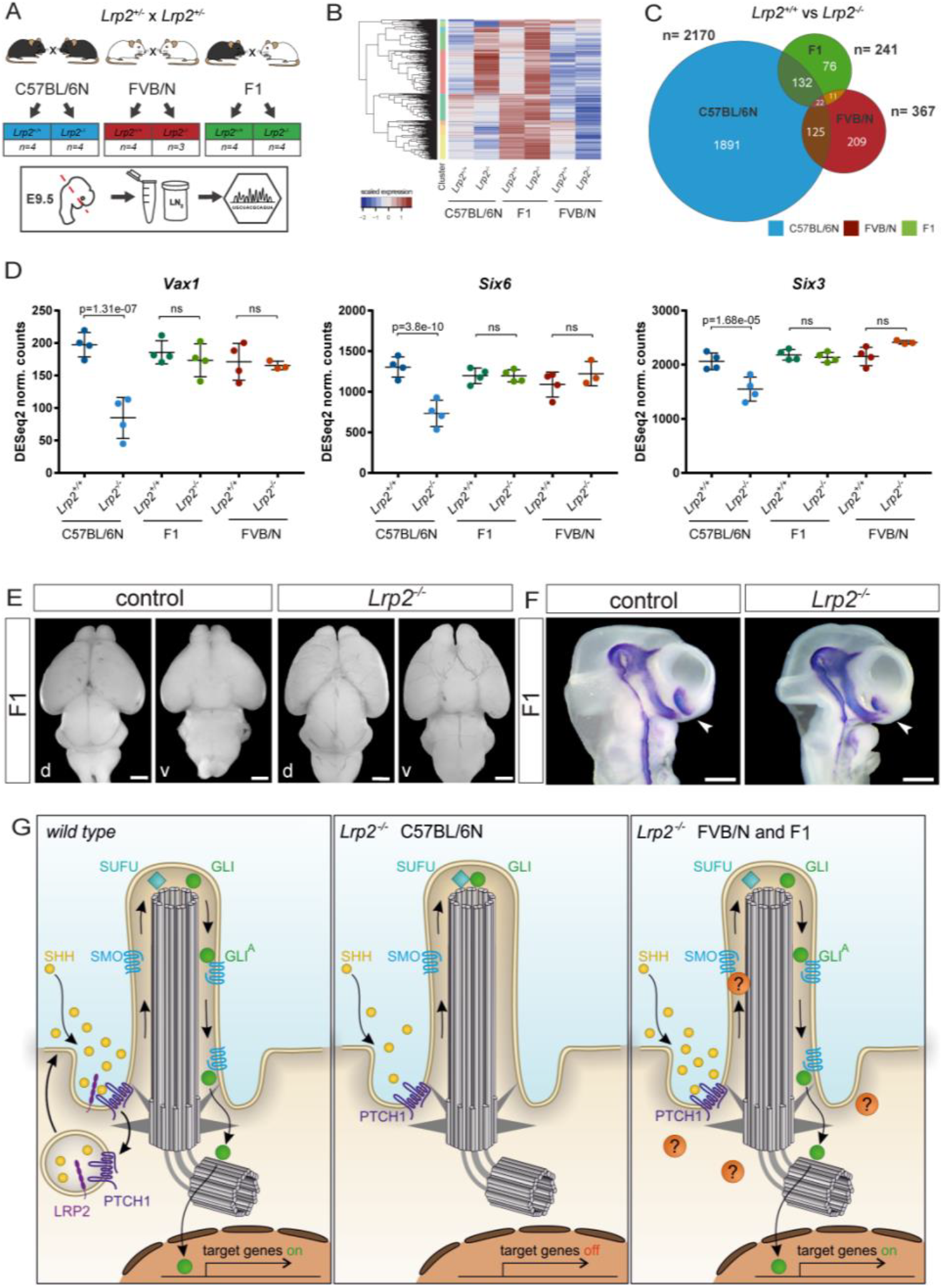
Dominant effect of FVB/N specific rescue modifier genes. (**A**) Collection of 24 somites staged *Lrp2*^+/+^ and *Lrp2*^−/−^ embryonic heads for RNA deep sequencing from C57BL/6N, FVB/N and F1 background. (**B**) Heatmap for RNA sequencing results. (**C**) Venn diagram with numbers of DEGs comparing *Lrp2*^+/+^ and *Lrp2*^−/−^ samples on C57BL/6N, FVB/N and F1 background. Illustrated are the numbers of distinct and overlapping DEGs. (**D**) DESeq2 normalized counts for *Vax1*, *Six6*, and *Six3*. (**E**) Brains from E18.5 control (refers to *Lrp2*^+/+^ and *Lrp2*^+/−^) (n = 17) and *Lrp2*^−/−^ F1 embryos (n = 19) in dorsal and ventral view. *Lrp2*^−/−^ F1 mice never displayed HPE (0/19) and showed normal separation of the forebrain hemispheres. Olfactory bulbs were present in *Lrp2*^−/−^ F1 brains. Scale bars: 1 mm. d: dorsal, v: ventral. (**F**) Whole mount ISH hybridization for *Shh* on E10.5 embryonic heads. Both, controls (*Lrp2*^+/+^ and *Lrp2*^+/−^) (n = 6) and *Lrp2*^−/−^ F1 embryos (n = 8) showed a typical *Shh* pattern including a prominent signal in the ventral telencephalon (arrowheads). Scale bars: 250 μm. (**G**) MODEL: Genetic modifiers rendering FVB/N and F1 more resilient to disturbances in the SHH pathway. Wild type: LRP2, a co-receptor for PTCH1, facilitates the internalization of SHH morphogen to activate the pathway. *Lrp2*^−/−^ C57BL/6N: SHH internalization and signaling are impaired. *Lrp2*^−/−^ FVB/N and F1: intact SHH pathway despite loss of LRP2, suggesting the presence of genetic modifiers. See also Figure S3.

Since none (0/68) of the hybrid C57BL/6N; FVB/N *Lrp2*^−/−^ F1 mutants suffered from HPE and SHH activity was fully rescued in *Lrp2*^−/−^ F1 embryos, we concluded that FVB/N allele-specific expression of yet unidentified genes has a dominant rescue effect. We hypothesize that such factors predispose neuroepithelial stem cells in FVB/N and F1 mice to maintain sufficient SHH activity and signaling despite the loss of LRP2 and thereby prevent HPE (Figure 3G).

### Identification of strain specific modifier genes involved in ciliogenesis and SHH pathway regulation

To identify strain specific modifier genes and pathways underlying HPE susceptibility in C57BL/6N, and HPE rescue in FVB/N and F1, we used the transcriptome data set described above (Supplementary Table 2). However, in this approach we focused on the strain specific transcriptome differences. Thus, we compared gene expression between mutant *Lrp2*^−/−^ samples on C57BL/6N, FVB/N and F1 backgrounds (Figure 4A - 4E) and most importantly between wild type (*Lrp2^+/+^*) samples comparing C57BL/6N, FVB/N and F1 backgrounds (Figure 4F - 4J). Venn diagrams show the number of DEGs in *Lrp2*^−/−^ embryos comparing the different backgrounds (Figure 4C) and in wild type *Lrp2^+/+^* embryos accordingly (Figure 4H). Given the dominant effect of the FVB/N background, we focused on DEGs, that were identified in both comparisons, C57BL/6N versus FVB/N as well as C57BL/6N versus F1 (Figure 4D and 4I and Supplementary Table 2). Importantly, out of the 1044 DEGs from the analysis comparing *Lrp2*^−/−^ mutant samples, 426 DEGs were also identified in the wild type *Lrp2^+/+^* comparisons (Supplementary Table 2). Thus, these DEGs, common between mutant and wild type comparisons of different strains, were not a consequence of LRP2 loss of function, but truly strain specific and therefore met our criteria of strain specific modifier candidates that can modulate signaling pathways in the wild type situation.

**Figure 4:**
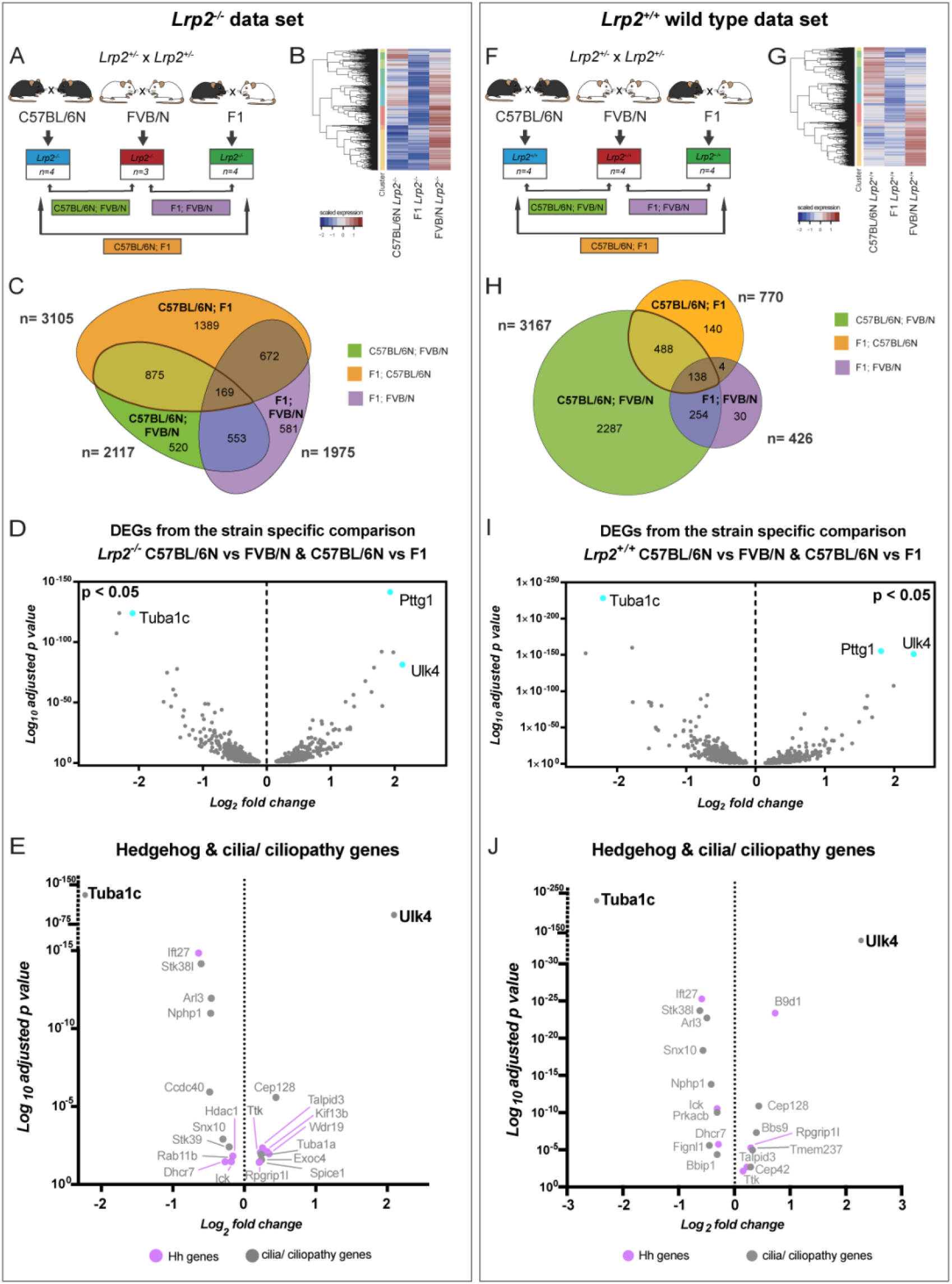
Comparative transcriptome analysis identifies strain-specific expression of genes involved in ciliogenesis and SHH pathway regulation. (**A**) and (**F**) RNA deep sequencing analysis on *Lrp2*^−/−^ (**A**) and *Lrp2*^+/+^ (**F**) embryonic heads on C57BL/6N, FVB/N, and F1 background. Comparison of the following expression profiles: C57BL/6N versus FVB/N, F1 versus FVB/N and C57BL/6N versus F1. (**B**) and (**G**) Pattern of the *Lrp2*^−/−^ and *Lrp2*^+/+^ heat maps for all different strains. (**C**) and (**H**) Venn Diagram (non-scaled) demonstrates the number of DEGs comparing *Lrp2*^−/−^ samples (**C**) and *Lrp2*^+/+^ samples (**H**). C57BL/6N versus FVB/N, F1 versus FVB/N and C57BL/6N versus F1 comparisons are demonstrated (colored ellipses and circles, respectively). Overlaps in two or three sets are indicated by the overlapping circles and by different color code. (**D**) and (**I**) Volcano plots showing DEGs for *Lrp2*^−/−^ samples (**D**) and *Lrp2*^+/+^ samples (**I**) that were identified in both comparisons, C57BL/6N versus FVB/N as well as C57BL/6N versus F1 hybrids. (**D**) 562 genes were significantly down- and 482 genes significantly up-regulated in *Lrp2*^−/−^ FVB/N and *Lrp2*^−/−^ F1 embryonic heads compared to *Lrp2*^−/−^ C57BL/6N samples. (**I**) 342 genes were significantly down- and 284 genes significantly up-regulated in *Lrp2*^+/+^ FVB/N and F1 samples compared to C57BL/6N heads. (**E**) and (**J**) Cilia, ciliopathy and hedgehog signaling pathway related genes after filtering DEGs from (**D**) and (**I**), shown in volcano plots for *Lrp2*^−/−^ (**E**) and *Lrp2*^+/+^ samples (**J**). Note, some of the cilia/ciliopathy genes are also hedgehog related genes and therefore labeled in purple.

Since there was a clear effect on the SHH pathway in the ventral forebrain organizer we next filtered those DEGs with a known function in the SHH pathway, cilia or ciliopathies from our strain specific data sets (Figure 4D and 4I and Supplementary Table 3). Hedgehog and cilia/ciliopathy relevant genes were selected according to published data (Breslow et al., 2018; Pusapati et al., 2018) which includes the Syscilia gold standard, and their expression was visualized by volcano plots (Figure 4E and 4J and Supplementary Table 3).

We determined that among cilia- and SHH-related genes 12 were down-regulated and 10 up-regulated in *Lrp2*^−/−^ FVB/N and F1 samples compared to C57BL/6N (Figure 4E). In the wild type *Lrp2^+/+^* comparison, we identified 20 differentially expressed cilia- and SHH-relevant genes, of which 11 were up- and 9 down-regulated in wild type FVB/N and F1 embryos compared to C57BL/6N (Figure 4J).

For functional analyses we selected top hits, based on the highest fold change and p-value identified in both, the wild type and mutant, comparisons (Figure 4E and 4J). Tubulin alpha 1C *(Tuba1c)* and Unc-51-like kinase 4 *(Ulk4)* were highly differentially expressed in a strain dependent manner.

*Tuba1c*, a component of tubulin, was expressed at five times lower levels in FVB/N *Lrp2*^−/−^ and wild type *Lrp2^+/+^* embryonic heads compared to both C57BL/6N genotypes. F1 samples showed also significantly lower expression levels than samples from the C57BL/6N strain (Figure S5A). *Tuba1c* is listed in the database of ciliary genes http://www.syscilia.org/goldstandard.shtml (van Dam et al., 2013; Pusapati et al., 2018), but data on its direct connection to the SHH pathway are very limited. However, pathway analysis has linked *Tuba1c* to the SHH “off” state (Fabregat et al., 2018; Rothfels, 2014).

*Ulk4*, which belongs to a family of serine/threonine kinases, has been shown to regulate acetylation of α-tubulin, an important post-translational modification of microtubules (Lang et al., 2016). *Ulk4* was 5 and 2 times higher expressed in FVB/N and F1 genotypes, respectively compared to C57BL/6N samples (Figure S5B).

### Ulk4 increases cellular SHH signaling capacity

We next analyzed the modifier genes in a context independent of LRP2 loss of function.

To test whether *Tuba1c* and *Ulk4* expression has functional effects on the SHH pathway we used a dual luciferase reporter assay (Christ et al., 2012; Sasaki et al., 1997; Taipale et al., 2000; Zhang et al., 2006) and quantified activity of SHH signaling upon expression of modifier candidates. Expression of *B9d1*, a SHH signaling modulator, was used as a positive control (Chih et al., 2012; Garcia-Gonzalo et al., 2011; Gerhardt et al., 2016). Overexpression of *B9d1* and *Ulk4,* respectively in transfected NIH-3T3 cells treated with SHH-Np resulted in a significant induction of SHH responsive and GLI driven relative luciferase levels compared to controls (Figure 5A). The induction was inhibited by KAAD-cyclopamine, showing a specific effect on the canonical SHH pathway. In contrast, cells transfected with *Tuba1c* showed no induction of GLI based luciferase activity after SHH stimulation compared to controls (Figure 5A).

**Figure 5:**
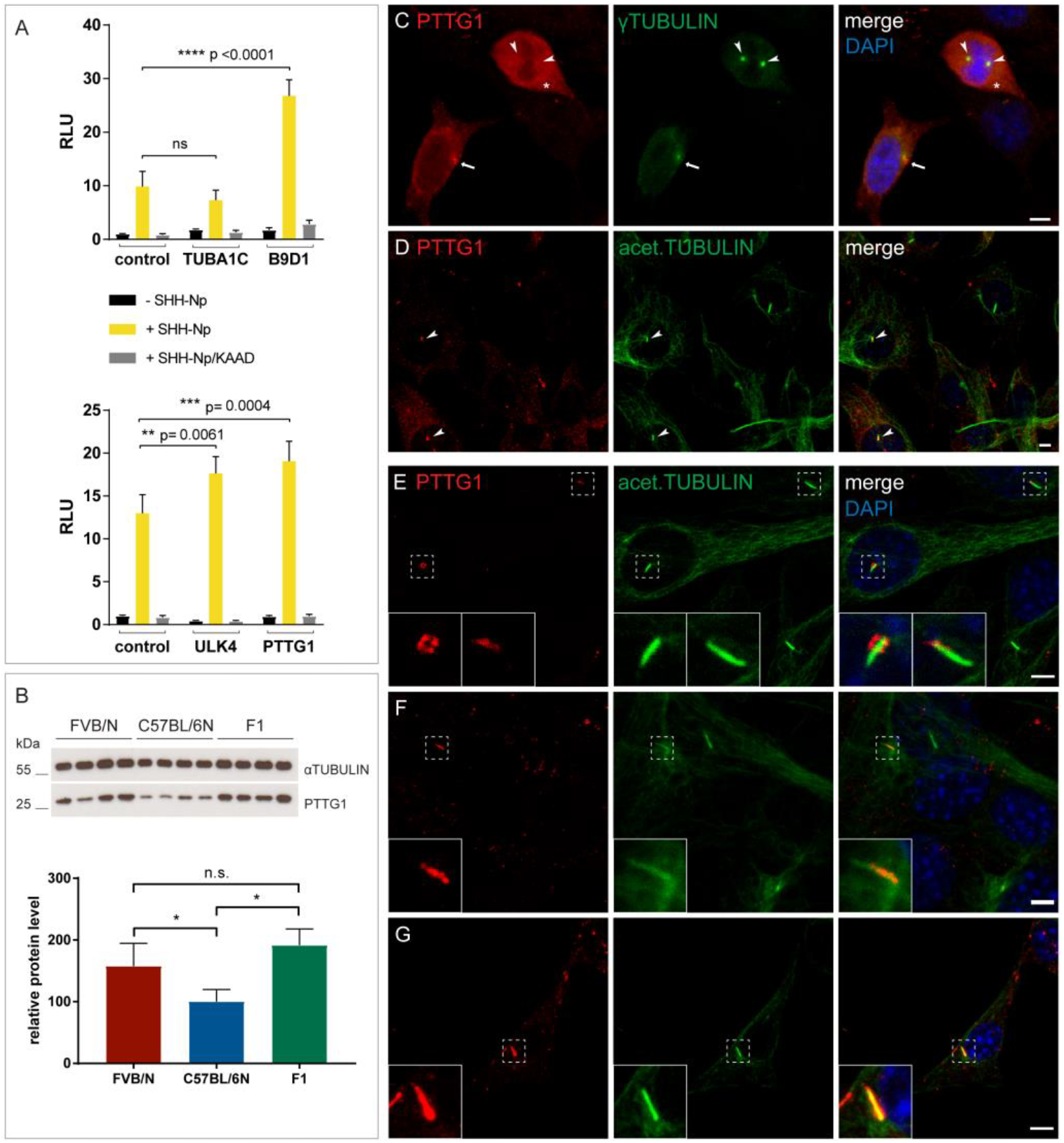
Novel SHH pathway modifier genes. (**A**) Luciferase reporter assay on NIH-3T3 cells. Significant induction of relative luciferase levels compared to controls, upon overexpression of *B9d1*, *Ulk4, and Pttg1*. *Tuba1*c expression showed no induction of SHH signaling. NIH-3T3 cells were treated with medium containing SHH-Np or control medium. KAAD-cyclopamine was used to confirm the canonical SHH pathway response. Significance assessed by two-way ANOVA; ** p < 0.001, *** p < 0.0001, ns: not significant; RLU: relative light units. (**B**) Western blot analysis of total PTTG1 protein levels in FVB/N, C57BL/6N, and F1 E8.5 *Lrp2*^+/+^ embryos (13 somites, n = 4 for each background). PTTG1 signals, normalized to α-tubulin, showed significantly higher PTTG1 levels in FVB/N and F1 compared to C57BL/6N embryos. Bar graph shows the quantification with significance assessed by un-paired t-test; * p < 0.05, ns: not significant. (**C**) - (**G**) Subcellular localization of PTTG1 in NIH-3T3 cells by immunocytochemistry. (**C**) Confocal microscopy detected PTTG1 in mitotic cells at the perinuclear region, concentrated at the centrosome and centrosomes (**C**, arrow and arrowheads) positive for γ-tubulin and in the cytoplasm (**C**, asterisk). (**D**) Interphase/quiescent cells showed PTTG1 localized to a subset of primary cilia stained with acetylated tubulin (**D**, arrowheads). Experiments were repeated at least five times in triplicates. Scale bars: 1 μm. (**E**) - (**G**) PTTG1 localized to a subset of primary cilia in a heterogeneous pattern. PTTG1 was detected in the periciliary region (**E**) and along the ciliary shaft, either at the proximal part (**F**) or covering entire cilium (**G**). Insets show the magnification of the chosen cilia (dashed line squares). Experiments were repeated at least five times in triplicates. Scale bars: 1 μm. See also Figure S5.

We conclude that *Ulk4*, which is highly expressed in the FVB/N and F1 rescue backgrounds, has a promoting effect on SHH signaling efficiency and represents a new SHH pathway modifier gene.

### Identification of PTTG1 as a novel SHH pathway modifier

We next focused on highly regulated genes that had not hitherto been associated with the primary cilium or SHH pathway and went back to the unbiased set of DEGs from all strain comparisons. The transcript with the highest fold change value and most significant p-value was pituitary tumor transforming gene 1(*Pttg1),* also known as securin (Figure 4D and 4I). Higher *Pttg1* transcript levels (Figure S5C) found in rescue backgrounds were reflected at the protein level. Western analysis detected significantly higher amounts of PTTG1 protein in FVB/N and F1 total embryos compared to C57BL/6N samples (Figure 5B). In addition, we demonstrated significantly higher immunofluorescence intensities for PTTG1 in the FVB/N and F1 forebrain neuroepithelium compared to C57BL/6N (Figure S5D).

We tested whether *Pttg1* can influence SHH signaling capacity. *Pttg1* overexpression in NIH-3T3 cells resulted in a significantly higher activation of the GLI luciferase reporter after SHH stimulation, compared to control vector transfection, and this was inhibited by KAAD-cyclopamine (Figure 5A). Our DESeq2 data and quantitative RT-PCR experiments showed a two-fold increase in *Gli1* levels in FVB/N rescue (Figure S5E and S5F) supporting the hypothesis that higher *Pttg1* expression levels could be one of the causative factors for more efficient SHH signaling.

### PTTG1 is a novel ciliary protein in the brain

We next analyzed the subcellular localization of PTTG1 and as expected (Moreno-Mateos et al., 2011; Tong et al., 2008) we detected PTTG1 in the perinuclear region concentrated at the centrosome and centrosomes of mitotic NIH-3T3 cells (Figures 5C, arrow and arrowheads respectively, and S5G: a). In some cells we also observed a dispersed PTTG1 signal in the cytoplasm (Figures 5C and S5G: a, asterisks). Interestingly, in non-mitotic NIH-3T3 cells we detected PTTG1 in the primary cilium, a microtubule-based organelle, essential for SHH signaling. PTTG1 localized to a subset of primary cilia (Figures 5D, arrowheads and S5G: b). We found PTTG1 in the periciliary region (Figure 5E) as well as along the shaft of the primary cilium (Figures 5F, 5G and S5G: b - c). Within the shaft PTTG1 protein localization was heterogenous, sometimes covering only the proximal region of the cilium (Figures 5F and S5G: b) and in other cases the entire ciliary shaft (Figures 5G and S5G: c). The variable localization of PTTG1 amongst different cilia suggested that it is not a structurally required component of the primary cilium, but may be shuttled into the shaft in a regulated fashion to support ciliary function.

We further analyzed if PTTG1 is also localized to cilia of the developing brain using cephalic explants from wild type embryos. In neuroepithelial stem cells from both mouse strains we detected PTTG1 localization to a subset of primary cilia (Figure 6A, arrowheads), in the periciliary region (Figure 6B: a, d) and in the shaft of the primary cilium labeled for ARL13b (Figures 6B: b, c, e, f and S6B: a’, a’’-b’, b’’).

**Figure 6:**
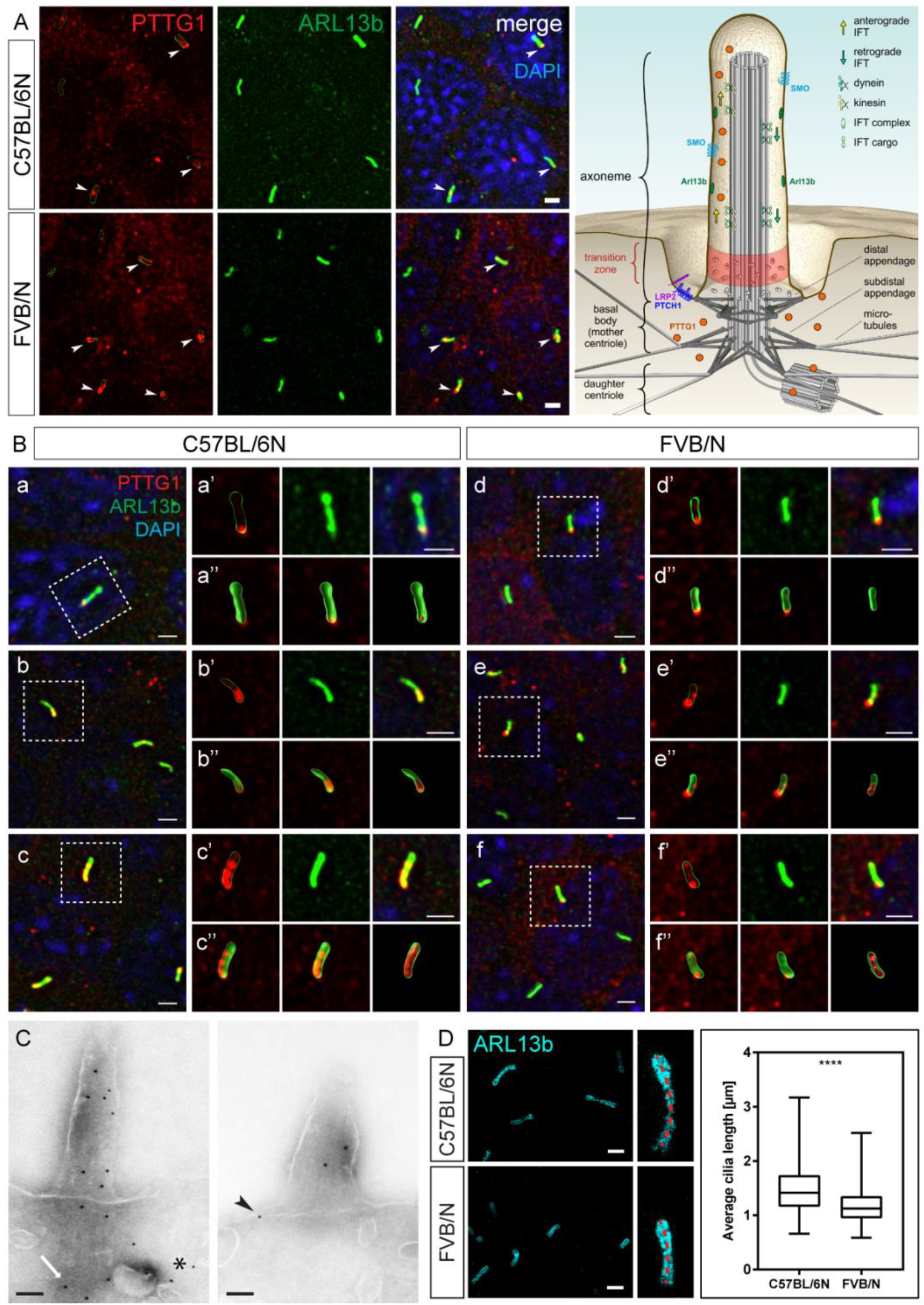
PTTG1 is a novel ciliary component in neuroepithelial stem cells. (**A**) Confocal microscopy on the E9.5 C57BL/6N (n = 6) and FVB/N (n = 6) cephalic explants shows PTTG1 in a subset of ARL13b positive primary cilia (arrowheads). Scale bars: 1 μm. (**B**) High resolution confocal 3D imaging: PTTG1 is present in the periciliary region (**a** and **d**) and in the ciliary shaft (**b**, **c** and **e**, **f**). Top panel insets (**a’**-**f’**) show magnified cilia (dashed squares) with single signals for PTTG1 (with outlining the cilium), ARL13b and merge, respectively. Bottom panel insets (**a’’**-**f’’**) show the same cilia as follows: reconstruction of ARL13b with non-reconstructed PTTG1 signal, optical section through this and reconstruction of both, ARL13b and PTTG1. Scale bars: 1 μm. (**C**) Immunogold labeling of PTTG1 in the forebrain neuroepithelium of C57BL/6N embryos at E9.5 showed clear localization of PTTG1 at the microtubule-based axoneme in the shaft of the primary cilium. PTTG1 was also localized to the daughter centriole (arrow) suggesting a role in microtubule stabilization and assembly of primary cilia. PTTG1 was also detected in the periciliary region (arrowhead) and few signals were detected in the cytoplasm (asterisk). Scale bars: 100 nm. (**D**) Differences in primary cilia length comparing C57BL/6N and FVB/N neuroepithelial stem cells. Primary cilia labeled for ARL13b were imaged in the anterior region of E9.5 cephalic explants using STED microscopy. Primary cilia of *Lrp2*^+/+^ FVB/N embryos (n = 7) were 20% shorter (1.19 μm ± 0.011 SEM; n = 1068) compared to cilia from *Lrp2^+/+^* C57BL/6N (n = 8) samples (1.49 μm ± 0.014 SEM; n = 908). Cilia length was measured as indicated by red dashed lines. The bar graph represents the mean cilia length with whiskers indicating minimal and maximal values and unpaired t-test for statistical analysis; **** p < 0.00001. Scale bars: 1 μm. (**E**) Schematic of the primary cilium with components, relevant to the present work. PTTG1 is a novel ciliary component localized to the periciliary region and the ciliary shaft. See also Figure S6.

Ultrastructural analysis revealed that PTTG1 was localized to the microtubule-based ciliary axoneme (Figure 6C, boxed and S6C) and to the periciliary region at the appendages (Figures 6C, arrowhead and S6C, arrowheads). PTTG1 localization at the base of and in the primary cilium is depicted in a graphical model (Figure 6E). Taken together our results identified PTTG1 as a novel microtubule-associated component of the primary cilium that can modulate efficiency of the canonical SHH signaling pathway.

### Cilia number and length differ between neuroepithelial stem cells from C57BL/6N and FVB/N wild type strains

Analyzing neuroepithelial stem cells from C57BL/6N and FVB/N cephalic explants revealed differences in cilia number per area (Figure S6B: a - b) and in cilia length between the strains (Figure 6D). Calculating the number of cilia in the anterior neural tube on the apical surface of the explants, we identified 645 cilia/mm^2^ surface area in FVB/N compared to 425 cilia/mm^2^ surface area in C57BL/6N explants (Figure S6B: a - b). We also used super resolution gated STED microscopy to image anterior neural tube explants from E9.5 C57BL/6N and FVB/N mice labeled for the regulatory GTPase ARL13b, which is localized to the membrane of the entire ciliary shaft. Strikingly, neuroepithelial stem cells of the rescue strain FVB/N had significantly shorter primary cilia compared to the HPE susceptible strain (Figure 6D).

## DISCUSSION

*Lrp2*^−/−^ mutant mice studied on a C57BL/6N, a mixed C57BL/6N; 129/SvEMS-Ter, CD1, and a C3H/HeNcrl background show fully penetrant forebrain defects and perinatal lethality (Sabatino et al., 2017; Spoelgen et al., 2005; Willnow et al., 1996). We and others previously found that a *Lrp2^267/267^* ENU mutant line on a predominantly FVB/N background survived to adulthood (Gajera et al., 2010; Zarbalis et al., 2004; Zywitza et al., 2018). These studies prompted us to perform a more rigorous study using a pure congenic *Lrp2*^−/−^ FVB/N line to analyze the molecular mechanisms underlying these dramatic rescue effects.

Here we demonstrate that the FVB/N background rescued HPE and underlying SHH signaling defects, as well as the previously described heart outflow tract phenotype of *Lrp2*^−/−^ C57BL/6N mice (Baardman et al., 2016; Li et al., 2015). Thus, genetic background strongly modifies the severity of congenital forebrain and heart defects in LRP2 deficient mice. To our knowledge, this is the only case where a HPE related gene mutation leads to a fully penetrant phenotype on one background compared to 100% rescue on a second inbred strain. Mice carrying mutations in HPE genes and showing classic HPE features can display a lower penetrance or a milder phenotype on a different background strain, but these were not all or none effects. CDON (cell adhesion molecule-related/down-regulated by oncogenes) is a positive regulator of the SHH pathway (Hong and Krauss, 2018; Hong et al., 2020; Zhang et al., 2006); *Cdon^−/−^* mice on a C57BL/6 background show HPE but only have microforms of HPE on a 129S6 strain. Background modifier studies in early embryonic development where the FVB/N strain was used, are rare and could yet shed important light on the etiology of neural tube defects as reviewed in (Leduc et al., 2017). Exencephaly in *Cecr2* mutant mice shows strain specific differences in penetrance comparing BALB/cCrl and FVB/N, with the latter being a rescue background. Whole genome linkage analysis revealed chromosome 19 as a modifier locus. In subsequent expression profiling of chromosome 19 by microarray analysis, the authors identified DEGs, including *Arhgap19,* which is expressed at lower levels in BALB/cCrl. A functional link between *Arhgap19* expression levels and the phenotype penetrance was not investigated (Banting et al., 2005; Davidson et al., 2007; Kooistra et al., 2011).

In our study we combined transcriptome and functional analyses to identify disease relevant modifier genes. Our RNA sequencing approach identified *Tuba1c* and *Ulk4* as the top down– and up-regulated DEGs, respectively, (Figure 4E and 4J). Functional data on *Tuba1c* regarding ciliogenesis or SHH pathway regulation are very limited. In the Reactome Pathway Knowledgebase *Tuba1c* is linked to the hedgehog (HH) “off” state suggesting that this tubulin plays a role as a negative regulator in the SHH pathway (Fabregat et al., 2018; Rothfels, 2014).

*Ulk4* is postulated to play an essential role in normal brain development and has been genetically linked to increased susceptibility to developing schizophrenia in humans (Lang et al., 2014). Mice with *Ulk4* gene defects show, amongst other phenotypes, hydrocephaly, dilated brain ventricles and ependymal motile ciliary defects (Liu et al., 2016); but ULK4 function has not been associated with the SHH pathway before. It is hypothesized that ULK4 regulates neuronal function by acetylation of α-tubulin, an important post-translational modification of microtubules (Lang et al., 2014, 2016). This is an interesting aspect since thereby both top hit DEGs, *Ulk4* and *Tuba1c*, are associated with microtubule nucleation.

Here, we demonstrate a potential new function for ULK4 as it can enhance canonical SHH signaling capacity in a heterologous system (Figure 5A). Further, we suggest that ULK4 is a novel factor that might regulate the penetrance of SHH related congenital disorders.

In an unbiased approach we asked whether the top regulated modifier candidate gene *Pttg1* could potentially also affect SHH activity. PTTG1 is a known substrate for the anaphase promoting complex (APC) and associates with separin until activation of the APC (Hagting et al., 2002; Mei et al., 2001; Thornton and Toczyski, 2003; Yanagida, 2000). Various functions for PTTG1 have been described including control of mitosis, DNA repair, transcriptional activity, and cell migration (Genkai et al., 2006; Havens and Walter, 2011; Hellmuth et al., 2020; Holt et al., 2008; Xiang et al., 2017; Yan et al., 2015; Zheng et al., 2015). The protein has tumorigenic activity and the gene is amplified in various human tumors of the breast, uterus, lung and thyroid ^74–78^. High expression of *Pttg1* has been also correlated with aggressive forms of brain tumors, including medulloblastoma and glioblastoma (Salehi et al., 2013; Yan et al., 2015). During embryonic development *Pttg1* expression has been reported in the human and murine brain (Boelaert et al., 2003; Karsten et al., 2003; Tarabykin et al., 2000). A recent study demonstrated a role of PTTG1 in microtubule nucleation (Moreno-Mateos et al., 2011). However, PTTG1 has so far not been associated with microtubule-based axoneme function of the primary cilium or with canonical SHH pathway modulation in the cilium. Here we show that *Pttg1* overexpression can indeed modulate SHH responsiveness in transfected cells (Figure 5A). Therefore, we conclude that PTTG1 is a new SHH pathway component and can promote canonical SHH signaling efficiency.

We also discovered a novel and seemingly dynamic localization of PTTG1 to the periciliary region and to microtubule-based axoneme of the cilium in quiescent/interphase NIH-3T3 and in neuroepithelial stem cells of the developing brain (Figures 5, S5, 6 and S6). The ciliary localization supports the hypothesis that PTTG1 modulates the SHH pathway via cilia-associated mechanisms. So far, only a relatively small number of regulatory proteins have been reported to be involved in both cytokinesis and ciliogenesis (Gromley et al., 2005; Kim et al., 2005; Pan et al., 2007; Park et al., 2008; Shah et al., 2008; Smith et al., 2011; Spektor et al., 2007; Vertii et al., 2015; Zuo et al., 2009). The anaphase promoting complex (APC), important for cytokinesis, is found to be also localized to the basal body of the primary cilium during interphase where its activity regulates disassembly of the primary cilium (Wang et al., 2014).

Our data provide evidence that PTTG1, besides its function as a cell cycle regulator associated with the centrosome, also plays a novel role in ciliogenesis and ciliary function, e.g. SHH signaling in interphase/quiescent cells. We propose that higher expression levels of PTTG1 predispose neuroepithelial stem cells in FVB/N mice to more efficient SHH signaling by enhancing microtubule repolymerization of the ciliary axoneme after cell division (Moreno-Mateos et al., 2011). Knockdown of PTTG1 has been shown to attenuate microtubule repolymerization after nocodazole treatment, which leads to microtubule depolymerization (Moreno-Mateos et al., 2011). Efficient microtubule repolymerization after cell division is crucial for assembly of the primary cilium and therefore SHH signaling (He et al., 2017). In our study, enhanced ciliogenesis is also supported by in average higher number of cilia per area in the FVB/N neuroepithelium compared to C57BL/6N.

Interestingly, all three top regulated modifier candidate genes, *Tuba1c*, *Ulk4*, and *Pttg1*, identified in our screen, are linked to microtubule function, suggesting that the expression profile of these genes in a FVB/N background provides an advantageous environment for the microtubule based ciliary function and SHH signaling, and ultimately a protective effect against developmental defects (Figure 7). These factors are most likely not necessary for establishing SHH signaling but have positive effects on the overall efficiency of the pathway. Combinatorial expression profiles of these candidate modifier genes, and likely other still uncharacterized modifier candidates, ensure sufficient SHH activity in the absence of LRP2 on a FVB/N background.

**Figure 7:**
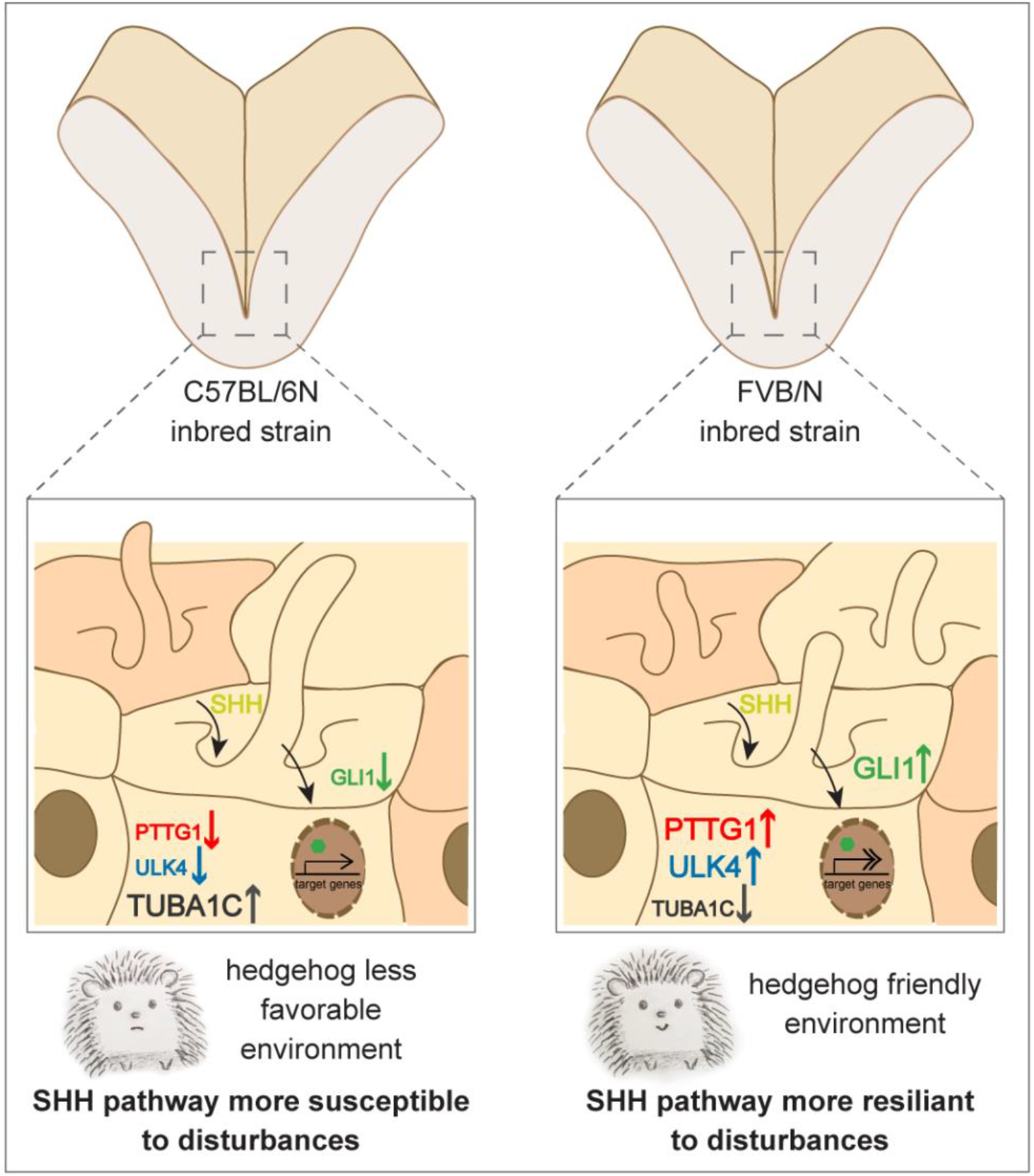
Proposed model. Sufficient SHH signaling is crucial for proper specification of the developing ventral neural tube and therefore essential for subsequent separation of the cortical hemispheres. Strain specific expression of candidate modifier genes leave the neuroepithelial stem cells of C57BL/6N embryos more susceptible to disturbances of the SHH pathway and therefore to congenital brain disorders such as HPE. FVB/N neuroepithelial stem cells with higher *Pttg1* and *Ulk4* and lower *Tuba1c* expression show enhanced SHH signaling and seem to be more resilient to disturbances of the SHH pathway. TUBA1c, ULK4, and PTTG1 are linked to microtubule function and PTTG1 is localized to the ciliary axoneme. The levels of the modifiers could enhance microtubule repolymerization of the ciliary axoneme after cell division and thereby facilitate SHH signaling. Further, in average shorter primary cilia could contribute to an overall more hedgehog friendly environment in FVB/N compared to C57BL/6N neural tissue.

Differences in ciliary factors between strains could be also reflected in different morphology of primary cilia. Indeed, our super resolution imaging revealed striking differences in the morphology of primary cilia between C57BL/6N and FVB/N neuroepithelial stem cells (Figures 6 and S6), suggesting that the genome of wild type mice from different mouse strains has a substantial influence on the shape of primary cilia. This could have functional consequences and relevant clinical implications considering the complex signaling function of the primary cilium (Gigante and Caspary; Goetz and Anderson, 2010; Nachury and Mick, 2019; Singla and Reiter, 2006). Little is known about the regulation of cilia length and the implication of variable cilia length in health and disease including ciliopathies (Fliegauf et al., 2007; Gerdes et al., 2009). Interactions of ciliary components, e.g. prominin with ARL13b, have been shown to play a role in maintenance of length of the primary cilia throughout the animal kingdom (Jászai et al., 2020). Studies in mammalian cells suggest that increased soluble tubulin production leads to longer cilia and cytosolic tubulin stabilization results in shorter cilia (Sharma et al., 2011; Wang and Dynlacht, 2018; Wang et al., 2013). Wang et al. showed that the complex formed by cell cycle regulator APC and its co-activator Cdc20 at the basal body maintains the optimal ciliary length and is crucial to shorten the cilia when cells exit from quiescent stage (Wang et al., 2014). Since PTTG1 is a substrate for APC (Mei et al., 2001; Yanagida, 2000), and also found at the base of the cilium in our study, this conceptually links the function of PTTG1 to cilia length regulation.

We conclude that the FVB/N inbred mouse strain has advantageous ciliary composition and morphology and identify plausible molecular candidates that render the strain more resilient to disturbances of the SHH pathway (Figure 7). Our study highlights the importance of mouse strain dependent penetrance of phenotypes. The expression profiles that differ profoundly between different wild type strains during early developmental stages provide a unique resource, that to our knowledge has not been available before. Considering the huge effects of the FVB/N background this resource is particularly valuable to understand genetic robustness in brain and heart development. Furthermore, we show that genetics in the mouse combined with functional cell biological studies can define novel molecular mechanisms that potentially underly variability in the penetrance of human cilia-associated neurodevelopmental disorders. Identification of disease relevant modifier genes in the mouse can provide important insights into the etiology and prevention of human congenital disorders.

## Supporting information

Supplemental spread sheet Table 2

Supplemental spread sheet Table 3

## ACKNOWLEDGEMENTS

We are grateful to Manfred Ströhmann for excellent work in the mouse husbandry. We thank Anke Scheer for excellent technical assistance. Wei Chen and Mirjam Feldkamp from the MDC genomics platform performed RNA deep sequencing. Alexandra Klaus-Bergmann provided helpful discussions on the study. Petra Schrade helped with scanning electron microscopy. Christina Schiel performed immunogold electron transmission microscopy. We thank Thomas Willnow for acquisition of financial support for the project. We would like to thank Gary Lewin for helpful discussions, intellectual input and critical reading of the manuscript. We are grateful to David Mick for critically reading the manuscript and intellectual input.

## AUTHOR CONTRIBUTION

N.M. conceived and performed experiments and analyzed data. I.K. conceived and performed experiments, analyzed data, and wrote parts of the manuscript. F.W. performed DESeq2 analysis and analyzed data, J.G. and A.L. performed experiments and analyzed data. H.G. and M.L. helped with STED microscopy and data analysis. M.R. and A.S. helped with confocal microscopy data analysis. B.P. carried out immunogold electron microscopy analysis. N.H. provided intellectual input, and edited the manuscript. A.H. conceptualized and supervised the study, secured funding and wrote the manuscript.

## METHODS

## KEY RESOURCES TABLE

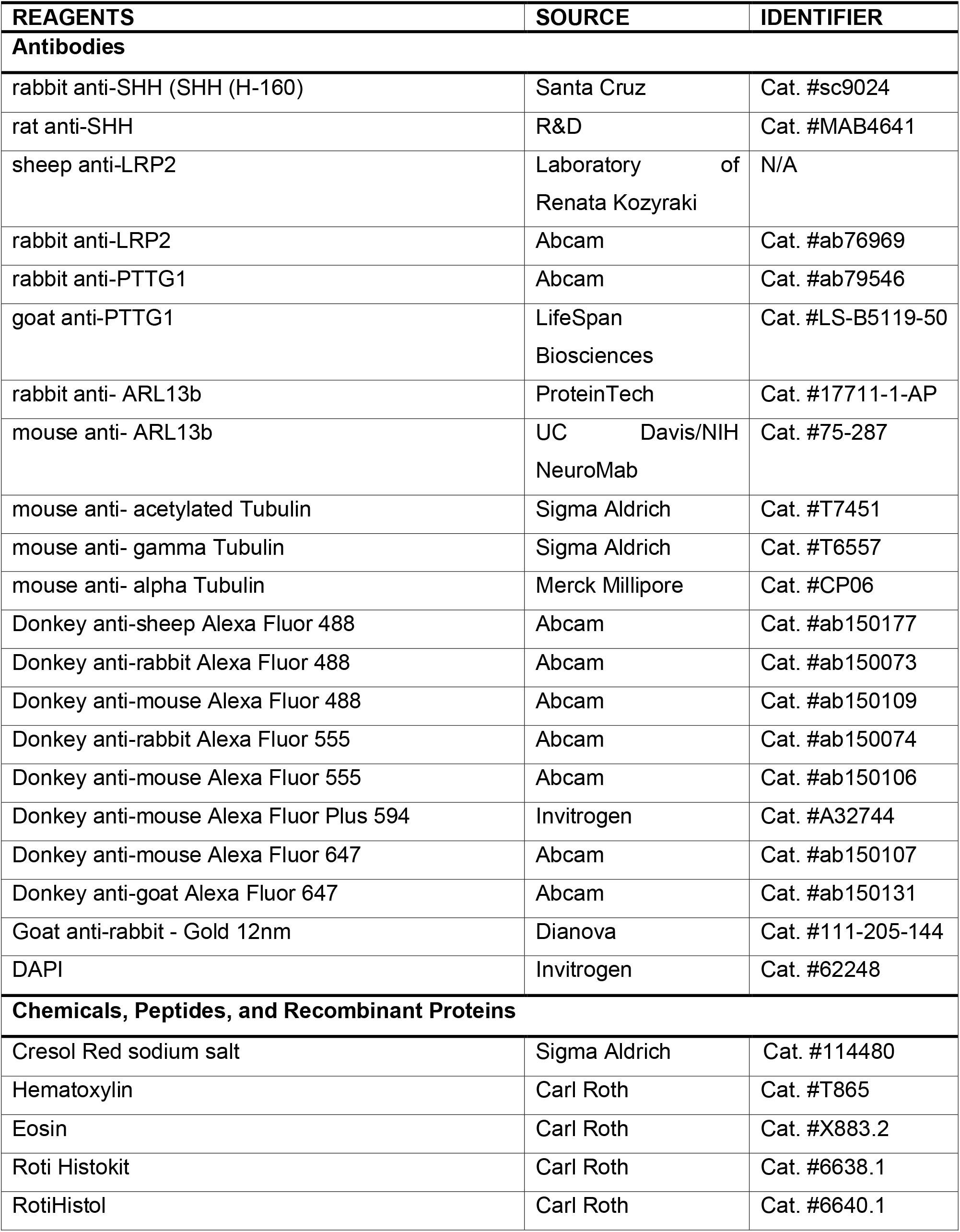
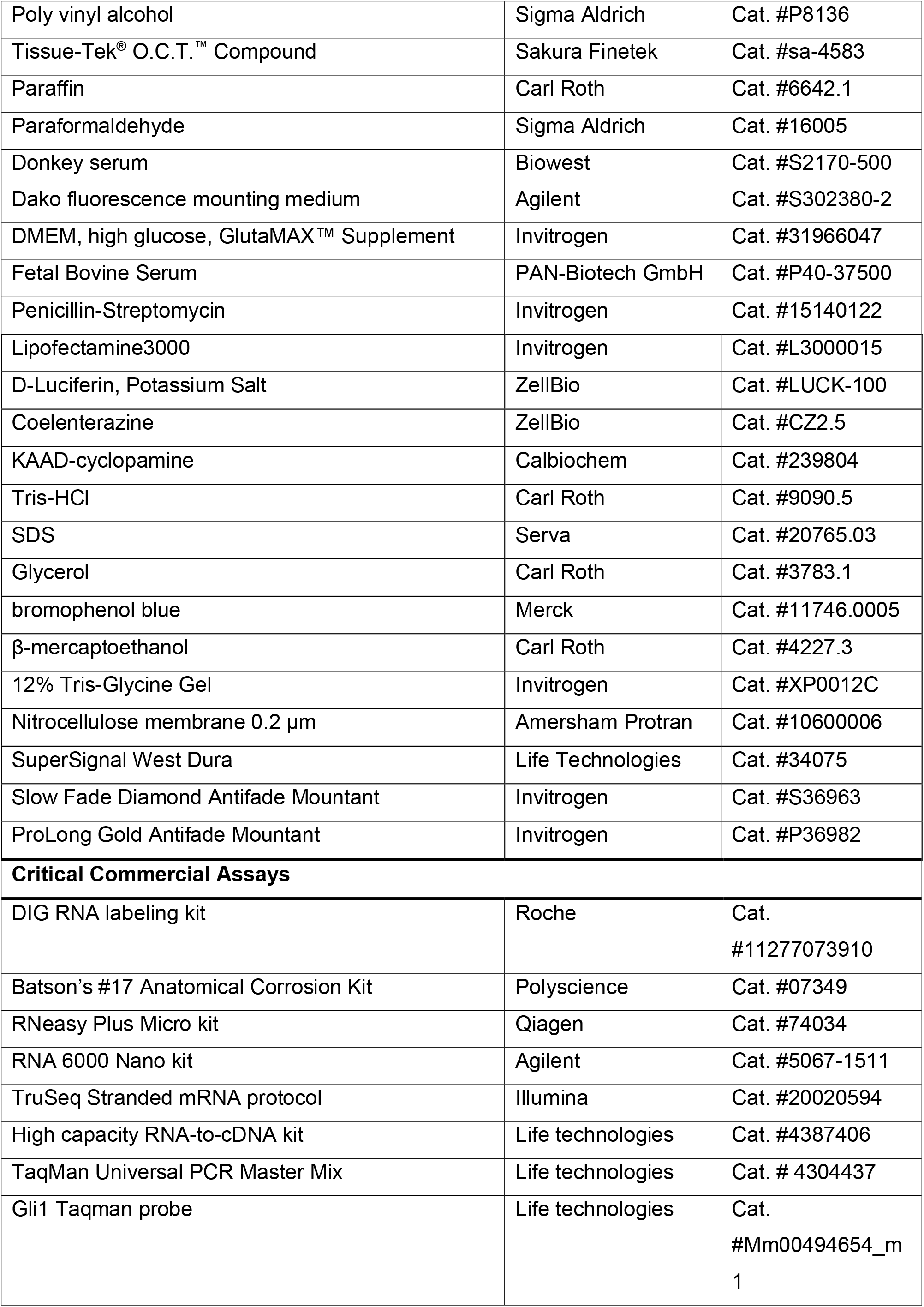
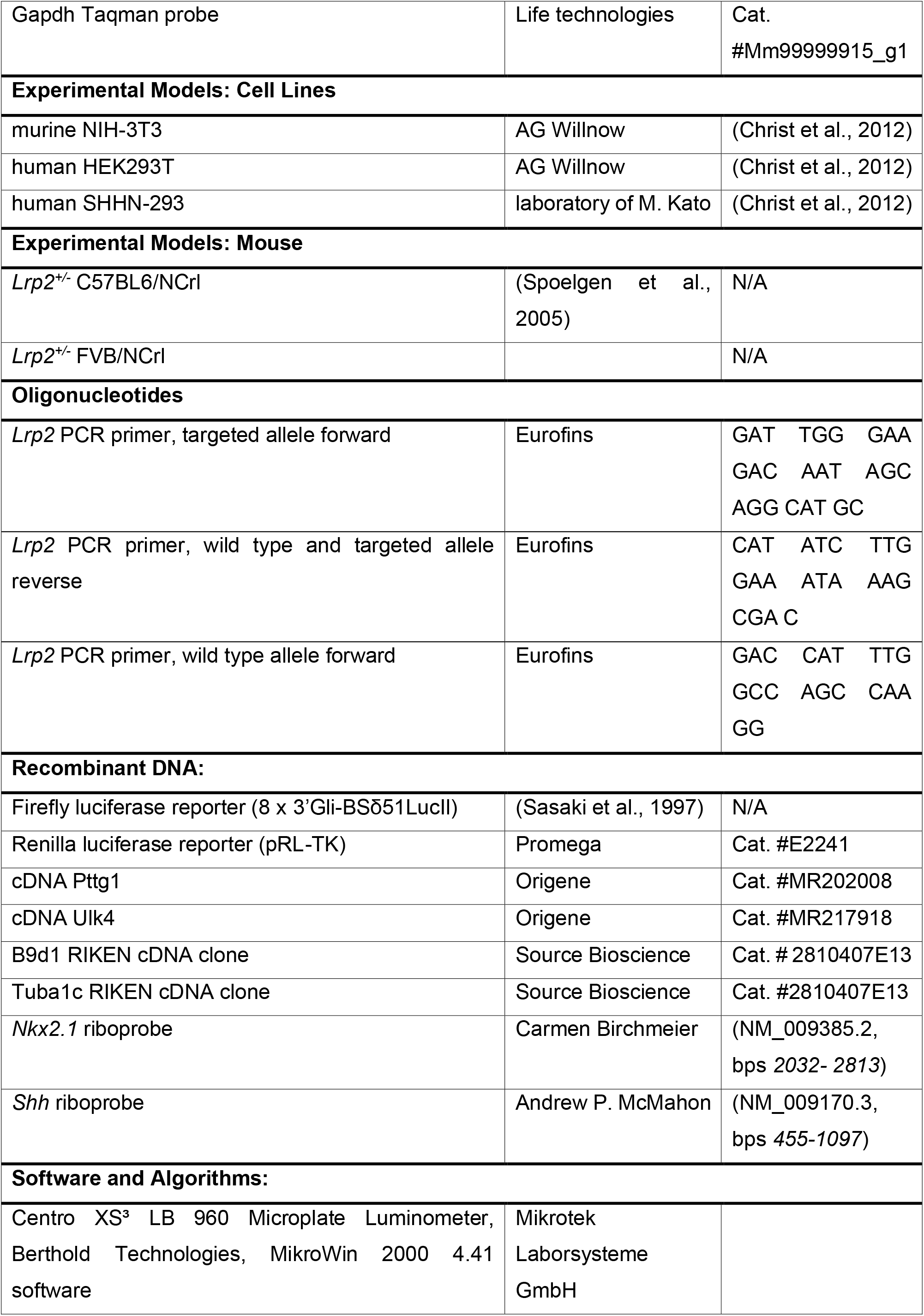
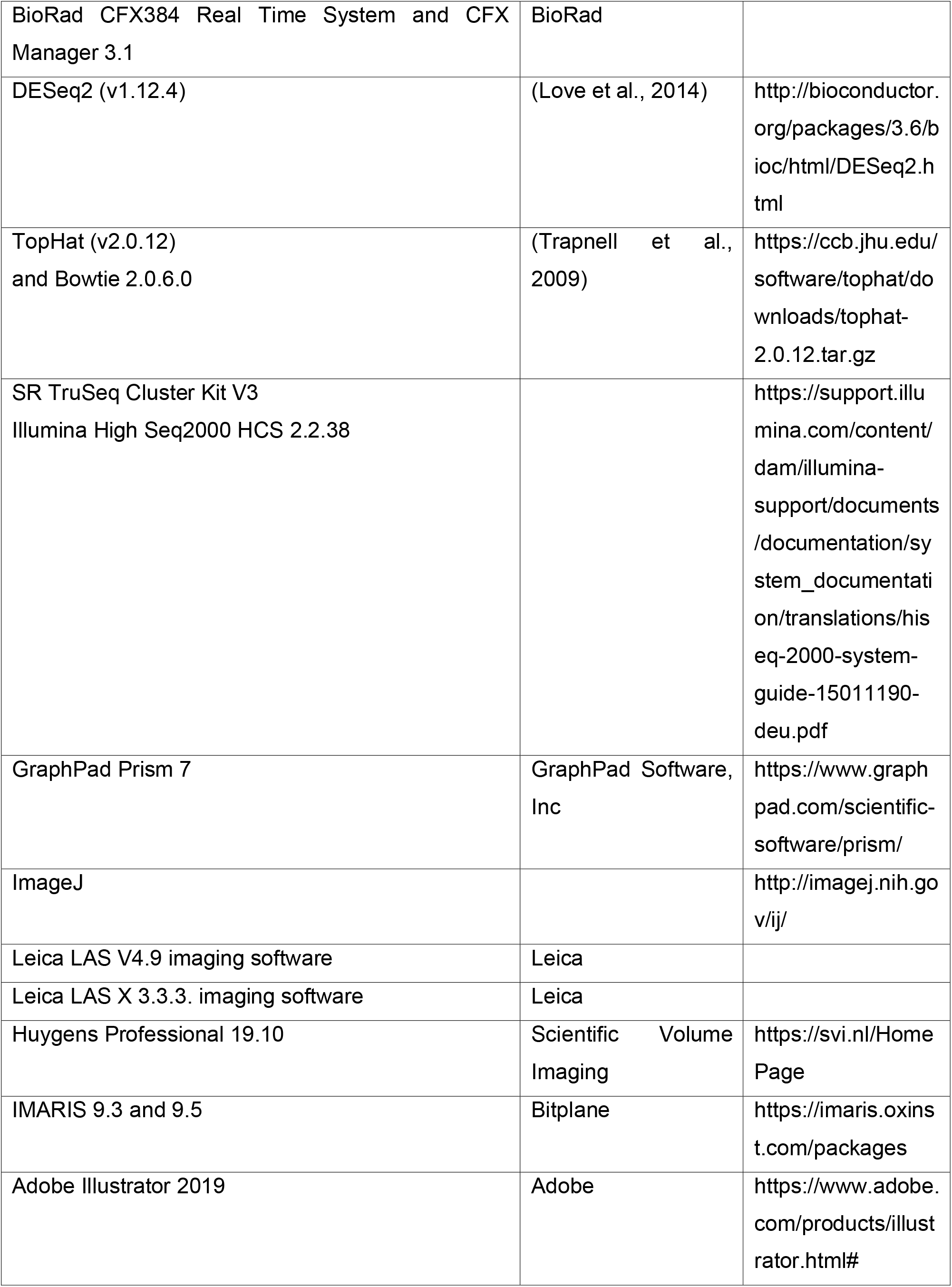
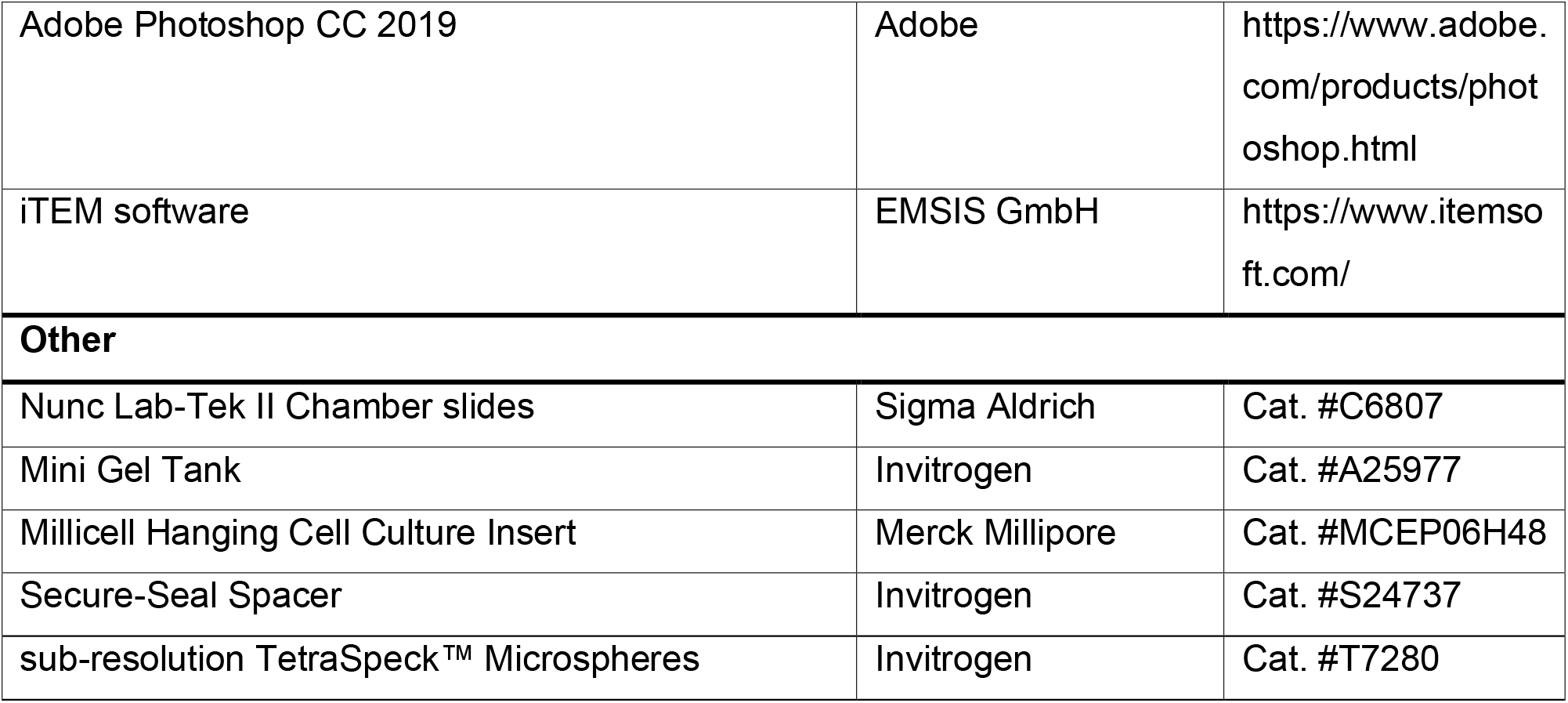

## RESOURCE AVAILABILITY

### Lead Contact

Further information and requests for resources and reagents should be directed to and will be fulfilled by the Lead Contact, Annette Hammes.

### Materials Availability

This study did not generate new unique reagents. *Lrp2*^+/−^ mouse line on FVB/N background will be available upon request.

### Data and Code Availability

The RNA sequencing datasets generated in this study have been made available as Supplementary Table 2 (excel file).

## EXPERIMENTAL MODEL AND SUBJECT DETAILS

### Animals

Experiments involving animals were performed according to institutional guidelines following approval by local authorities (X9005/12). Mice were housed in a 12 hours light-dark cycle with ad libitum food and water. The generation of mice with targeted disruption of the *Lrp2* gene has been described before (Willnow et al., 1996). The *Lrp2* mutant mouse line was crossed onto a pure C57BL/6NCrl background in our laboratory (Spoelgen et al., 2005) (herein referred to as *Lrp2* mutant C57BL/6N. For this study the *Lrp2*^+/−^ C57BL/6N line was backcrossed for more than 12 generations to obtain a congenic mouse line on a FVB/NCrl background (herein referred to as *Lrp2* mutant on FVB/N background).

We used mice > 8 weeks of age for timed matings for our experiments. Analyses of the congenital defects were carried out in *Lrp2*^−/−^ and in somite matched *Lrp2^+/+^* and/or *Lrp2*^+/−^ control littermates on a C57BL/6NCrl and FVB/NCrl background. For the generation of the F1 hybrid *Lrp2* mutant embryos, *Lrp2*^+/−^ FVB/NCrl females were crossed with *Lrp2*^+/−^ C57BL/6NCrl males. PCR genotyping was performed from yolk sac genomic DNA for E8.5 embryos and from tail biopsies for older embryos.

## METHOD DETAILS

### Histology

Standard NISSL and Hematoxylin & Eosin staining were performed on paraffin sections. Embryos were fixed in 4% paraformaldehyde in PBS at 4°C, overnight, embedded in paraffin and cut with 10 μm thickness. NISSL staining was performed in 0.1% Cresol Red solution (Sigma Aldrich, Cat. #114480) for 3-10 minutes. Hematoxylin & Eosin staining was performed according to manufacturer’s instructions with incubation times of 5 min in Hematoxylin (Carl Roth, Cat. #T865) and 3 min in Eosin (Carl Roth, Cat. #X883.2). Sections were dehydrated and mounted in Roti Histokit (Carl Roth, Cat. #6638.1). Staining was visualized using the Leica DM5000B microscope using the LAS-X 3.3.3. Software.

### *In situ* Hybridization

Whole mount *in situ* hybridization (WISH) was carried out as described previously (Hammes et al., 2001). *In situ* hybridization (ISH) on sections was performed as described in (Jensen and Wallace, 1997), except that the signal was enhanced by performing the color reaction in the presence of 10% poly vinyl alcohol (Sigma Aldrich, Cat. #P8136). Probe synthesis was conducted with the components of the DIG RNA labeling kit (Roche, Cat. #11277073910). Plasmids for generating *in situ* probes were generated from the following mRNA sequences: *Nkx2.1* (NM_009385.3, bps *2032 - 2813*) kindly provided by Carmen Birchmeier, *Shh* (NM_009170.3, bps 455 - 1097) kindly provided by Andrew P. McMahon (University of Southern California).

### Injection of heart samples with Batson’s pigment

Polymeric dye injections were used to visualize the ascending aorta and the pulmonary artery in isolated E18.5 mouse hearts applying the Batson’s #17 Anatomical Corrosion Kit (Polyscience, Cat. #07349). Batson’s #17 Blue pigment was added to Base Solution A in the amount of 2%, mixed vigorously and divided into two equal parts. 24 ml of the Catalyst was added to the first 100 ml of Base Solution A/pigment mix. 24 drops of Promoter C were carefully added to the second half of the Base Solution A/pigment and mixed slowly. Both solutions were mixed together and stirred. After both parts were mixed it took 30 to 45 minutes until the solution was fully polymerized and until then the experiment should be finished. The same procedure was repeated for Batson’s #17 Red pigment. Injections of the blue and red pigment solution into the right and left ventricle, respectively, were made with a disposable polyethylene syringe and 23G needle under a stereomicroscope (Leica MZ 10F). Images of the heart were taken after the injection (Leica LAS V4.9 imaging software).

### RNA library generation and sequencing

For the generation of an RNA library embryonic heads of E9.5 embryos were dissected, snap frozen and stored at −80°C until RNA isolation. RNA was extracted using RNeasy Plus Micro kit (Qiagen, Cat. #74034), checked on Bioanalyzer (Agilent RNA 6000 Nano kit, Cat. #5067-1511), and samples with RIN>9 were used to prepare the cDNA library. Library preparation for mRNA sequencing was performed according to the Illumina SR TruSeq Stranded mRNA protocol (Cluster Kit V3, Illumina, Cat. #20020594) on 23 embryonic head RNA samples from mice with 24 somites. Sequencing was performed on the Illumina HiSeq2000 system with HCS 2.2.38 software.

Sequencing reads were aligned to a SNP-infused (FVB/NCrl and C57BL/6NCrl SNPs, accordingly) mouse genome (Ensembl GRCm38.77) using TopHat v2.0.12 with Bowtie 2.0.6.0. The number of reads that mapped to a gene was counted using the HTseq-count v0.6.0 with default parameters. All expressed genes (n = 10,861) have been used for differential expression analysis using DESeq2 v1.12.4 (Love et al., 2014). We defined genes as differentially expressed meeting our significance threshold of FDR ≤0.05. Allele specific expression was determined by counting the reads matching the C57BL/6N or FVB/N genotype in the F1 mRNA-seq data.

### Differential gene expression analysis

mRNA-seq quantifications were derived from exon-mapped, paired-end reads. Expression quantification was followed by read normalization, size factor estimation and differential expression analysis using DESeq2 v1.12.4 (Love et al., 2014).

For this analysis we included all genes that we consider to be expressed, defined as having at least 100 reads in 23 out of 26 samples (n = 10,861). We considered a gene differentially expressed when it met genome-wide significance thresholds of FDR ≤ 0.05. We performed three different comparisons: a) genotype comparison, where we assessed expression differences for each mouse strain separately (C57BL/6N, FVB/N or the F1 strain) comparing *Lrp2*^−/−^ versus *Lrp2^+/+^* samples; b) strain comparison, where the different mouse strains, C57BL/6N versus FVB/N and F1, were compared in *Lrp2^+/+^* and *Lrp2*^−/−^ samples; and c) the interaction of genotype and strain effects, where the interplay of strain and LRP2-deficiency was assessed to understand strain differences that were observed between *Lrp2^+/+^* versus *Lrp2*^−/−^ samples.

### Real-Time Quantitative Reverse Transcription PCR (Real-Time qRT-PCR)

Total RNA from E9.5 embryos heads was isolated and cDNA was synthesized by High capacity RNA-to-cDNA kit (Life technologies, Cat. #4387406). Quantitative PCR was performed using TaqMan Universal PCR Master Mix (Life technologies, Cat. #4304437) with the BioRad CFX384 Real Time System used on a BioRad C1000 Touch Thermal Cycler. The expression of *Gli1* (Taqman probe, Life Technologies, Cat. #Mm00494654_m1) was normalized to *Gapdh* (Life Technologies, Cat. #Mm99999915_g1). Transcript levels relative to *Gapdh* were calculated using the deltaCt method. Data were analyzed in GraphPad Prism 7 (GraphPad Software, Inc) using an unpaired t-test.

### Cell Culture

The NIH-3T3 and HEK293T cell lines were obtained from T. Willnow (MDC) (Christ et al., 2012). SHHN-293 cells were originally kindly provided by M. Kato (Stanford School of Medicine). Cells were maintained in DMEM (Invitrogen, Cat. #31966047) with 10% Fetal Bovine Serum (FCS, PAN-Biotech GmbH, Cat. #P40-37500) and 1% Penicillin-Streptomycin (Invitrogen, Cat. #15140122). Cells were transfected with Lipofectamine™ 3000 Transfection Reagent (Invitrogen, Cat. #L3000015) according to the manufacturer’s instructions.

The following overexpression constructs were used: cDNA Pttg1 (Origene, Cat. #MR202008), cDNA Ulk4 (Origene, Cat. #MR217918), B9d1 RIKEN cDNA clone (Source Bioscience, Cat. #0710008G05), Tuba1c RIKEN cDNA clone (Source Bioscience, Cat. #2810407E13).

### Dual luciferase reporter assay

NIH-3T3 cells, seeded in 24-well plates, were transiently co-transfected with the gene of interest or proper empty plasmid, respectively, GLI-dependent firefly luciferase reporter (8 x 3’Gli-BSδ51LucIl) (Sasaki et al., 1997) and a constitutive Renilla luciferase reporter (pRL-TK; Promega, Cat. #E2241) at a ratio 10:10:1 using Lipofectamine3000, according to manufacturer’s protocol.

24 h later the medium was replaced with conditioned medium from HEK293 cells stably secreting SHH-Np (SHHN-293 cells; kindly provided M. Kato, Stanford School of Medicine) or control medium from parental HEK293T cells, at a 1:10 dilution, in medium containing 0.5% FCS. For control studies KAAD-cyclopamine (Calbiochem, Cat. #239804) at a 50 nM concentration or solvent were added to the conditioned medium with SHH-Np.

After 48 hours of stimulation, activity was assayed with D-luciferin (ZellBio GmbH, Cat. #LUCK-100) and results were normalized to the corresponding Renilla activity, assayed with coelenterazine (ZellBio GmbH, Cat. #CZ2.5). Software for Dual Luciferase reporter assay measurements: Centro XS³ LB 960 Microplate Luminometer, Berthold Technologies, MikroWin 2000 4.41 software from Mikrotek Laborsysteme GmbH. Experiments were performed in triplicates, referring to three different wells assayed the same day, in minimum of 3 independent experiments, and results are shown as mean and standard deviation. Statistical analyses were performed using Two-way ANOVA with Dunnett’s multiple comparisons test.

### Western Blot Analysis

E8.5 whole embryos were lysed in SDS lysis buffer (60 mM Tris-HCl pH-6,8, 2% SDS, 10% glycerol, 0.01% bromophenol blue, 1.25% beta-mercaptoethanol). Samples were heated to 95°C for 10 min, centrifuged for 2 min at 13,200 rpm and stored at −20°C until used. Equal amounts of samples were subjected to a 12% Tris-Glycine Gel (Invitrogen, Cat. #XP0012C) in a Mini Gel Tank (Invitrogen, Cat. #A25977). The resolved proteins were transferred onto a nitrocellulose membrane (Amersham Protran 0.2 μm, Cat. #10600006) using a wet electroblotting system (Bio-Rad Mini Protean II Cell) followed by immunoblotting. 5% non-fat dry milk in TBS-T (0.1% Tween-20) was used for blocking at room temperature for 1 hr. Primary antibodies were used overnight at 4°C as follows: rabbit anti-PTTG1 (Abcam, Cat. #ab79546, 1:5000), mouse anti-alpha Tubulin (Merck Millipore, Cat. #CP06, 1:2000), Signal was detected by SuperSignal West Dura (Life Technologies, Cat. #34075) with Otimax 2010 X-Ray Film Processor (PROTECT) and the result was quantified using ImageJ, with unpaired t-test as a statistical analysis.

## SAMPLE PREPARATION FOR CONFOCAL MICROSCOPY

### Immunofluorescence on sections

Standard immunofluorescence was performed on cryo- and paraffin sections. For cryosections, PFA fixed embryos were infiltrated with 15% and 30% sucrose in PBS for up to 24 hours depending on the stage, embedded in O.C.T. (Tissue-Tek^®^ O.C.T., Sakura Finetek, Cat. #sa-4583) and cut into 10 μm coronal sections. For paraffin sections PFA fixed embryos were dehydrated, incubated in RotiHistol (Carl Roth, Cat. #6640.1), embedded in paraffin and cut into 10 μm sections. Standard immunohistochemical analysis was carried out by incubation of tissue sections with primary antibodies at the following dilutions: rabbit anti-SHH (Santa Cruz, Cat. #sc9024, 1:50), sheep anti-LRP2 antiserum (1:4000) kindly provided by the Laboratory of Renata Kozyraki, rabbit anti-LRP2 (Abcam, Cat. #ab76969, 1:1000), rabbit anti-PTTG1 (Abcam, Cat. #ab79546, 1:100). Bound primary antibodies were visualized using secondary antisera conjugated with Alexa Fluor 488, 555, 647 (1:500, Abcam). All samples were counterstained with DAPI (Invitrogen, Cat. #62248. Sections were mounted with Dako Fluorescence Mounting Medium (Agilent, Cat. #S302380-2).

### Immunocytochemistry

For immunocytochemistry NIH-3T3 cells were seeded in chamber slides (Sigma, Cat. #C6807) in regular 10%FCS/DMEM medium conditions. After 24 h cells were rinsed with PBS(1x) fixed with 4% PFA for 15 min, RT and permeabilized with PBS-TritonX 0.25% for 20 min. Blocking with 10% donkey serum/PBS-TritonX for 1 hr, was followed by a standard incubation with primary and secondary antibodies as described above, with dilutions as follows: rabbit anti-PTTG1 (Abcam, Cat. #ab79546, 1:100), goat anti-PTTG1 (LifeSpan Biosciences, Cat. #LS-B5119-50, 1:100), mouse anti-ARL13b (UC Davis/NIH NeuroMab, Cat. #75-287, 1:500), rabbit anti-ARL13b (ProteinTech, Cat. #17711-1-AP, 1:1500), mouse anti-acetylated Tubulin (Sigma, Cat. #T7451, 1:1000), mouse anti-γTubulin (Sigma, Cat. #T6557, 1:200). Walls from the chambers were disassembled according to manufacturer’s instruction and samples were mounted with Dako Fluorescence mounting medium. Each experiment was repeated at least five times n triplicates.

### Preparation of mouse cephalic explants for immunofluorescence analysis

Explants were prepared analogically as described in (Echevarria et al., 2002). E9.5 mouse embryos were collected and the neural tube was cut open along the dorsal midline, from caudal to rostral direction, using an insect needle. The neural folds were precisely cut above the heart and placed on the sterile filter (Millipore, Cat. #MCEP06H48) on the petri dish in the drop of PBS(1x). The floor plate at the level of the cephalic flexure was pinched in order to unfold the tissue with the ventricular part facing up. Filter was placed in the 6-well plate containing DMEM/10% FCS and explants were incubated at 37°C, with 5% CO2 and 95% humidity for 3-4 hours to flatten and recover. The explants were washed gently in PBS(1x), fixed 1 hr in 4% PFA, and subjected to the standard immunofluorescence protocol described above, with 2x O/N for primary antibody incubation, in the dilutions as follows: rabbit anti-PTTG1 (Abcam, Cat. #ab79546, 1:100), goat anti-PTTG1 (LifeSpan Biosciences, Cat. #LS-B5119-50, 1:100), mouse anti-ARL13b (UC Davis/NIH NeuroMab, Cat. #75-287, 1:500). Secondary antibodies conjugated with Alexa Fluor 488 were used to visualize primary cilia and with Alexa Fluor 647 were used to visualize PTTG1 signals. Explants were flat-mounted in Slow Fade Diamond Antifade Mountant (Invitrogen, Cat. #S36963) using Secure-Seal Spacer (Invitrogen, Cat. #S24737) in order to match the refractive index (RI) of the mountant and the immersion media of the glycerol objective, and to prevent distortions of the explant by pressure, shrinking and hardening of the mountant.

## CONFOCAL MICROSCOPY IMAGE AQUISITION

Image acquisitions of tissue sections were carried out using either a Leica SPE or Leica TCS SP8 confocal microscope using a HC Pl Apo 20× NA 0.75 MultiIMM and HC Pl Apo 63× NA 1.3 oil immersion objective. All samples that were compared either for qualitative or quantitative analysis were imaged under identical settings for laser power, detector and pixel size. Cells were imaged with a Leica TCS SP5 confocal microscope equipped with a ACS Apo 63× oil NA 1.3 immersion objective.

The forebrain region of the explant samples was imaged en face with a Leica TCS SP8 confocal microscope using a HC Pl Apo 63× NA 1.3 glycerol immersion objective with a WD of 0.3 mm to enable high-resolution imaging with minimal spherical aberrations of the thick specimen. Three regions per explants were imaged for n = 6 C57BL/6N and n = 6 FVB/N. High-resolution z-stack (60 nm pixel size, 12 bit, 0.2 μm z-step size) images of apical side of neuroepithelium were acquired with a z-piezo stepper. In all samples, Alexa Fluor 488 was excited by a 488 nm laser, detection at 500 - 550 nm, Alexa Fluor 555 was excited by a 555 nm laser, detection at 570 - 620 nm, Alexa Fluor 647 was excited by a 633 nm or 647 nm laser, detection at 660 - 730 nm and DAPI was excited at 405 nm, detection at 420 - 450 nm with a pinhole set to 1 AU.

## IMAGE PROCESSING AND ANALYSIS

### 3D Data processing, deconvolution and correction

Confocal Z-stacks of explants were subjected to a background correction and processed by deconvolution with the CMLE algorithm in order to obtain an improved signal to noise ratio and axial and spatial resolution using Huygens Professional 19.10 software (Scientific Volume Imaging). For optimal deconvolution the experimental PSF was calculated with Huygens PSF Distiller by using sub-resolution TetraSpeck™ Microspheres, 0.2 μm (Invitrogen, Cat. #T7280) embedded and acquired with same imaging conditions as the explant sample. The same deconvolved beads sample was used to estimate a chromatic aberration correction matrix (in x/y/z) which was used to correct the experimental sample data. To get isotropic voxel values the aspect ratio of the deconvolved bead sample was changed. This new sampling value was applied to all image data with the IMARIS Software before doing further segmentation and analysis steps.

### Immunofluorescence signal localization analysis

Localization of the protein of interest (PTTG1) within the primary cilium of mouse neuroepithelium was assessed using IMARIS software (Imaris 9.3 and 9.5, Bitplane) after applying raw data corrections (deconvolution, chromatic shift correction, non-isotropic imaging correction). 3D rendering and surfaces reconstruction of individual cilia was performed for two channels – one for cilia marker (ARL13b) and one for PTTG1 – and the classical object-based colocalization approach (individual segmentation and voxel based colocalization) with IMARIS XT module was used. To visualize the precise spatial coexistence of the two proteins in individual cilia, views from different angles were created and transparent LUTs as well as clipping planes were used to cut the 3D surfaces open to enable seeing inside cilia. Total number of cilia in each sample was assessed during the same processing steps from the reconstructed surface of ARL13b signal, and was given as number of cilia per area (Figure S6a).

### Quantification of PTTG1 immunofluorescence signal intensity

Z-stack images of the coronal sections were analyzed using ImageJ (Fiji, NIH). For the quantification of PTTG1 signals in the neuroepithelium the full Z-stack was used. Region of interest (ROI) was manually outlined as shown in the (Figure S5d) and the mean fluorescent intensity was measured with the ROI manager. The average intensity for each animal was used in the final quantification and unpaired t-test statistical analysis was performed to assess the significance.

### Immunofluorescence analysis of primary cilia using STED microscopy imaging

En face single-color stimulated emission depletion (STED) microscopy imaging was performed on the mouse cephalic explants, which were prepared as described above, with exception to the highly cross-absorbed secondary antibody Alexa Fluor Plus 594 (Invitrogen, Cat. #A32744) and ProLong Gold Antifade mountant (Invitrogen, Cat. #P36934) that were used to obtain optimal resolution. STED microscopy images were taken with a Leica SP8 TCS STED microscope (Leica Microsystems) equipped with a pulsed white-light excitation laser (WLL; ∼80 ps pulse width, 80 MHz repetition rate (NKT Photonics) and two STED laser for depletion at 592 nm and 775 nm. The system was controlled by the Leica LAS X software. Single-color STED imaging was performed by exciting Alexa Fluor Plus 594 at 594 nm and the depletion of its emission with the 775 nm STED laser. Time gated detection was set from 0.3 – 6 ns. The fluorescence signal was detected from 604 – 650 nm by a hybrid detector (HyD) at appropriate spectral regions separated from the STED laser. Images were acquired with a HC PL APO CS2 100×/1.40 NA oil objective (Leica Microsystems), a scanning format of 1,024 × 1,024 pixels, 8-bit sampling, 16x line averaging and 6x optical zoom, yielding a pixel size of 18.9 × 18.9 nm. Additionally, to every STED image, a confocal image with the same settings but just 1x line averaging was acquired.

All STED microscope images of the cilia within each independent experiment were acquired with equal settings. Single optical plane images were subjected to the cilia length quantification in ImageJ. Regions of interest (ROIs) were manually selected with the segmented line tool and the total lengths of single cilia were measured. The average cilia lengths for both mouse lines were checked for the statistical significance using unpaired t-test.

### Scanning electron microscopy

For scanning electron microscopy E8.5 embryos were dissected and fixed in 0.1 M sodium cacodylate buffer (pH 7.3/7.4) containing 2.5% glutaraldehyde. After rinsing in cacodylate buffer a postfixation step in 2% OsO_4_ for two hours was followed. Samples were dehydrated in graded ethanol series, osmicated, dried in critical point apparatus Polaron 3000, coated with gold/palladium MED 020 (BAL-TEC) and examined at the Zeiss scanning electron microscope (Gemini DSM 982).

### Transmission electron microscopy and immunogold labeling

The head region of mouse embryos at E9.5 was fixed with 3% freshly prepared formaldehyde/ 0.05% glutaraldehyde (EM-grade) in 0.1 M phosphate buffer for 1 hour at RT. After washing, samples were infiltrated with 2.3 M sucrose over night at 4°C and frozen. Semithin sections were prepared to identify the region of the neuroepithelium. Ultrathin cryosections according to Tokuyasu were labeled with anti-PTTG1 antibody (Abcam, Cat. #ab79546, diluted 1:50) and 12 nm colloidal gold anti-rabbit secondary antibody (Dianova, Cat. #111-205-144). Sections were contrasted and stabilized with a mixture of 3% tungstosilicic acid hydrate and 2.5% polyvinyl alcohol. EM pictures were taken at 80 kV with a Morgagni electron microscope (Thermo Fisher), equipped with a Morada camera and the iTEM software (EMSIS GmbH, Münster, Germany).

### Quantification and statistical analysis

Tests used to analyze the data were done using Prism 7 software (GraphPad) and are mentioned in the respective Figure legends. Figures were prepared using Adobe Illustrator 2019 software.

The term significant was used, if the p value of a result was smaller than 0.05. Exact p values, n numbers and biological replicates are reported in the Figure legends, supplementary tables, or according methods.

