## Supplemental Figures and Legends for "Identification of novel disease relevant genetic modifiers affecting the SHH pathway in the developing brain"

**SUPPLEMENTAL ITEMS:**

| | Normal outflow tract 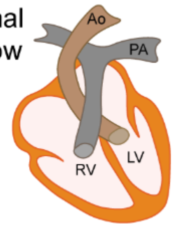 | DORV 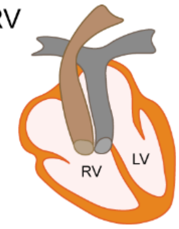 | CAT 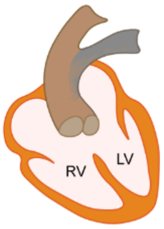 |
|---|---|---|---|
| C57BL/6N control | 7 | 0 | 0 |
| C57BL/6N *Lrp2*$^{-/-}$ | 0 | 1 | 15 |
| FVB/N control | 11 | 0 | 0 |
| FVB/N *Lrp2*$^{-/-}$ | 11 | 2 | 0 |
| F1 control | 9 | 0 | 0 |
| F1 *Lrp2*$^{-/-}$ | 5 | 4 | 0 |

**Supplementary Table 1: Frequency of congenital heart defects in *Lrp2*$^{-/-}$ embryos and controls on different strain backgrounds. Related to Figure 1 and Supplementary Figure 3**

The table indicates the number of embryos analyzed for each genotype and the numbers on phenotype penetrance.

CAT: common arterial trunk; DORV: double outlet right ventricle

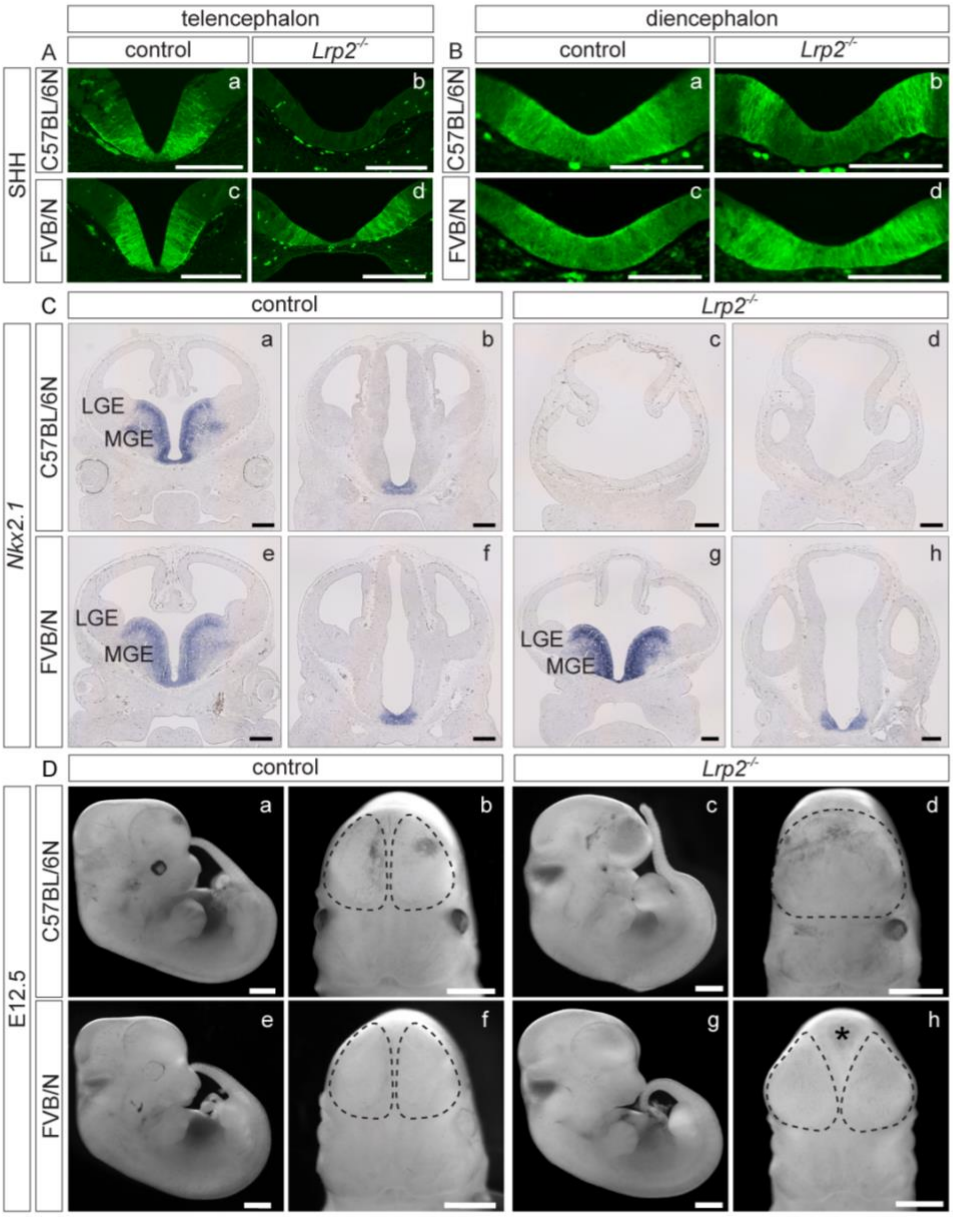


**Figure S2: Loss of ventral forebrain SHH and Nkx2.1 in *Lrp2$^{-/-}$* C57BL/6N mice leading to HPE is rescued in mutants on a FVB/N background. Related to Figure 2**

(**A**) - (**B**) Immunohistological detection of SHH on coronal sections of the forebrain in control (refers to *Lrp2$^{+/+}$* and *Lrp2$^{+/-}$*) and somite matched *Lrp2$^{-/-}$* embryos at E10.5.

(**A**) SHH protein was lost in the ventral telencephalon of *Lrp2$^{-/-}$* C57BL/6N embryos (**b**), (n = 3), compared to controls (**a**), (n = 5). *Lrp2$^{-/-}$* mice on a FVB/N background displayed normal SHH protein localization in the ventral telencephalon (**d**), (n = 6), comparable to controls (**c**), (n = 7). Scale bars: 250 µm.

(**B**) SHH in the ventral midline of the diencephalon was shifted to more lateral domains in *Lrp2$^{-/-}$* C57BL/6N embryos (**b**), (n = 3), compared to controls (**a**), (n = 5). *Lrp2$^{-/-}$*

FVB/N embryos presented with a normal SHH pattern (**d**), (n = 6), similar to controls (**c**), (n = 7). Scale bars: 100 µm.

(**C**) In situ hybridization (ISH) of coronal forebrain paraffin sections demonstrated loss of *Nkx2.1* expression at E12.5 in the medial ganglionic eminence (MGE) in *Lrp2*$^{-/-}$ C57BL/6N embryos (**c**, **d**), (n = 3) compared to controls (**a**, **b**), (n = 3). In contrast, all *Lrp2*$^{-/-}$ FVB/N embryos displayed normal expression of the transcription factor (**g**, **h**), (n = 3) comparable to controls (n = 2) in the basal telencephalon (**e**, **f**).

MGE: medial ganglionic eminence; LGE: lateral ganglionic eminence. Scale bars: 250 µm.

(**D**) Phenotypic characterization of control embryos and *Lrp2*$^{-/-}$ mutants on a C57BL/6N and a FVB/N background at E12.5, presented in a lateral view and of embryonic heads presented in a frontal view. 100% of *Lrp2*$^{-/-}$ C57BL/6N embryos (n = 21) displayed severe forebrain malformations, small eyes (**c**) and a single forebrain hemisphere (**d**, dashed line circle) compared to controls (**a**, **b**), (n = 31). 100% of *Lrp2*$^{-/-}$ FVB/N embryos (n = 52) showed normal appearance in the lateral view (**g**) comparable to controls (**e**), (n = 102). They developed separated forebrain hemispheres (**h**, dashed line circles), highlighted in the frontal view. Scale bars: 1 mm.

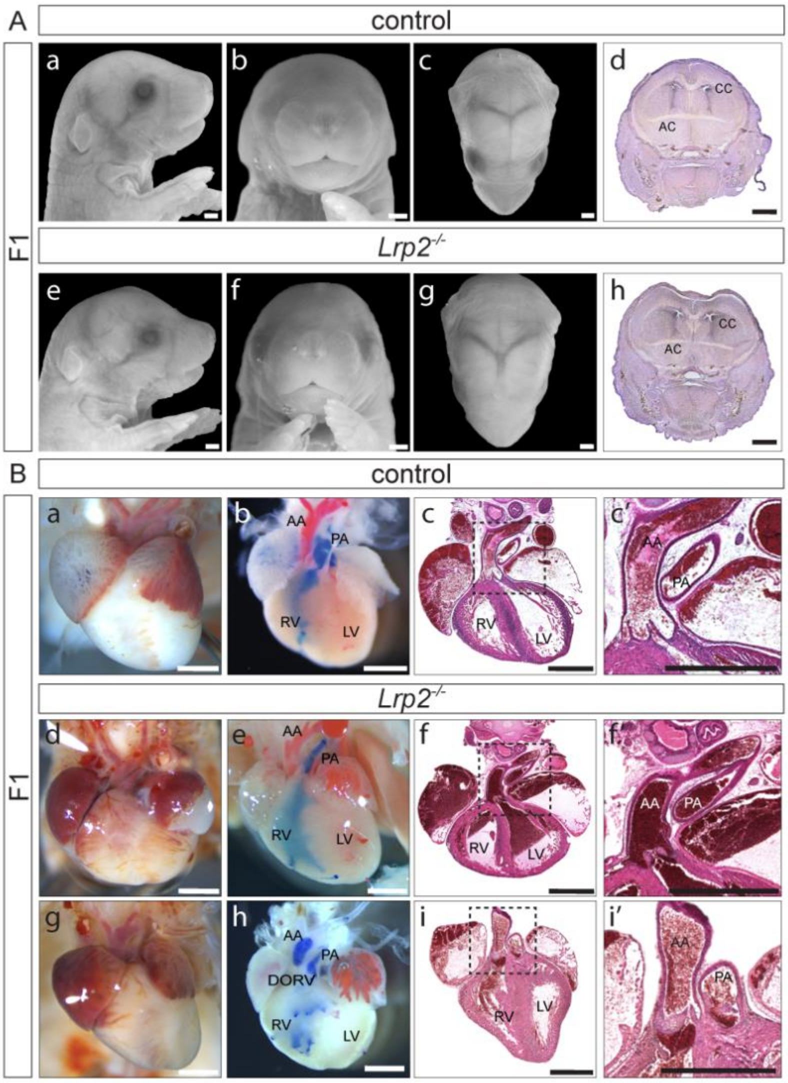


**Figure S3: Rescue of congenital defects in *Lrp2$^{-/-}$* mutants on a hybrid F1 (C57BL/6N; FVB/N) background. Related to Figure 3**

(**A**) E18.5 embryonic heads shown in the sagittal (**a** and **e**), frontal (**b** and **f**) and dorsal (**c** and **g**) view. All *Lrp2$^{-/-}$* F1 mutants (n = 49) showed normal craniofacial features compared to controls (*Lrp2$^{+/+}$* and *Lrp2$^{+/-}$*; n = 78). NISSL stained coronal cryosections of E18.5 *Lrp2$^{-/-}$* heads on a F1 background (**h**), (n = 17) showed normally separated ventricles, normal corpus callosum (CC) as well as anterior commissure (AC) structures comparable to controls (**d**), (n = 16). Scale bars: 1 mm.

(**B**) F1 control (**a**), (n = 3) and F1 *Lrp2$^{-/-}$* (**d** and **g**), (n = 4) embryonic hearts at E18.5. Batson's red pigment was injected into the left ventricle, followed by injection of blue pigment into the right ventricle (**b**, **e**, **h**). Frontal plane of H&E stained paraffin sections through hearts of control embryos (**c**, **c'**), (n = 6) and *Lrp2$^{-/-}$* F1 embryos (**f**, **f'**, **i**, **i'**), (n = 5) demonstrated the outflow tract anatomy. 55.6 % (5/9) of *Lrp2$^{-/-}$* F1 embryos showed normal heart morphology and outflow tract anatomy (**d** - **f**) compared to controls (**a** - **c**). 44.4% *Lpr2*$^{-/-}$ F1 mice (4/9) showed a double outlet right ventricle (DORV) (**g** - **i**). Importantly, a common arterial trunk (CAT) phenotype was never observed in any of the *Lrp2* F1 mutants (0/27). This suggests a partial rescue of the heart outflow defects in *Lrp2* F1 mutants compared to *Lrp2* mutants on a C57BL/6N background (see **Figure S1**: **C**, **D**, **F** and **Supplementary Table 1**). Scale bars: 1 mm. AA: ascending aorta; PA: pulmonary artery; LV: left ventricle; RV: right ventricle

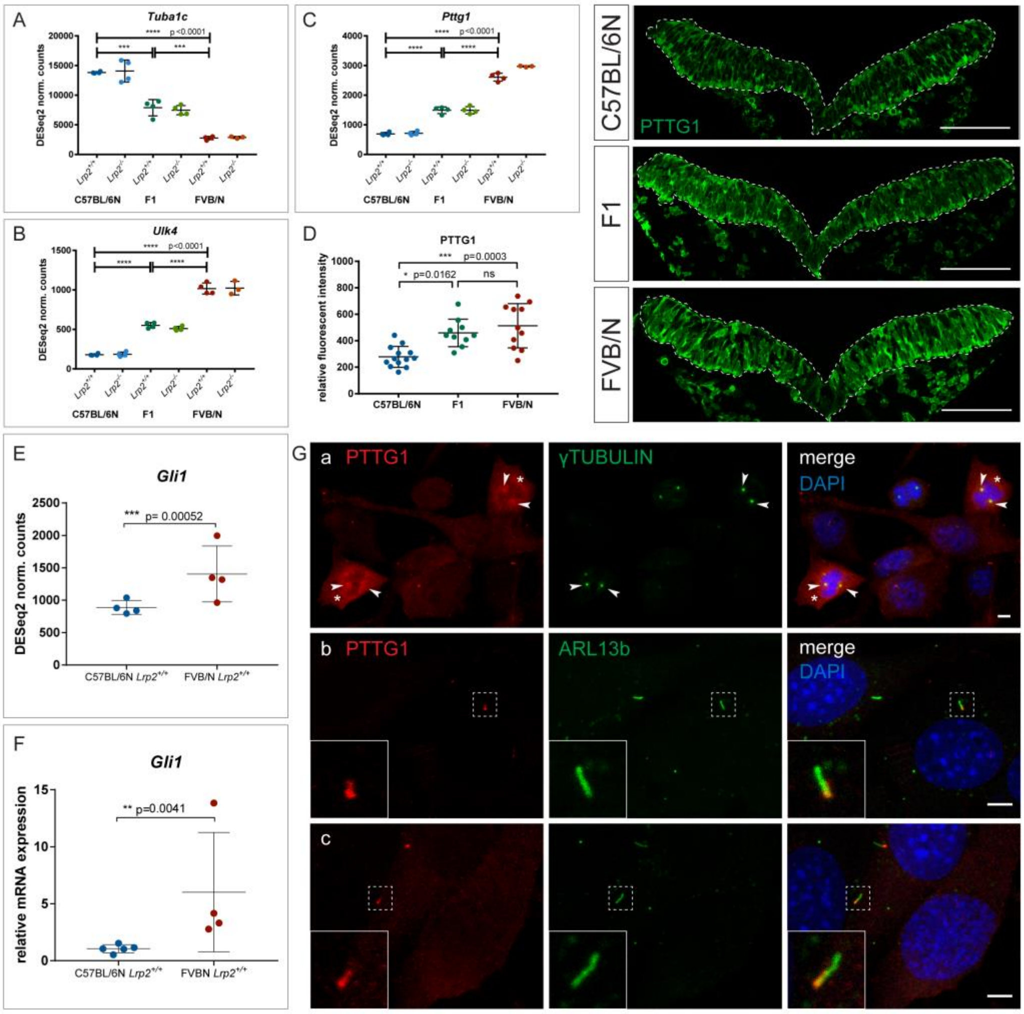


**Figure S5: Candidate modifier gene expression and PTTG1 localization in cell culture and *in vivo.* Related to Figure 5**

(**A – C**) DESeq2 normalized counts for *Tuba1c, Ulk4 and Pttg1*. *Tuba1c* showed significantly lower expression levels for F1 and FVB/N backgrounds, compared to C57BL/6N. *Ulk4* and *Pttg1* showed significantly higher expression levels in embryonic head samples of both genotypes on a FVB/N and F1 compared to C57BL/6N background.

(**D**) Immunofluorescence intensity of PTTG1 protein was measured in the neuroepithelium (dashed line) from E8.5 C57BL/6N (n = 5), F1 (n = 3) and FVB/N (n = 4) control embryos (*Lrp2*$^{+/+}$ and *Lrp2*$^{+/-}$). A total of 3 coronal sections from each embryo (10 and 11 somites) were examined. Significantly higher PTTG1 protein levels were found in the FVB/N and F1 embryos compared to C57BL/6N. Significance assessed by unpaired t-test and represented on the graph, * $p < 0.05$, *** $p < 0.0001$. Scale bars: 100 µm.

(**E**) DESeq2 normalized counts for *Gli1* showed significantly higher expression level for *Lrp2*$^{+/+}$ FVB/N compared to *Lrp2*$^{+/+}$ C57BL/6N samples.

(**F**) Relative mRNA expression level analysis by qRT-PCR confirmed significantly higher *Gli1* expression for *Lrp2*$^{+/+}$ FVB/N compared to *Lrp2*$^{+/+}$ C57BL/6N in E9.5 embryonic heads. *Lrp2*$^{+/+}$ C57BL/6N: n = 5; *Lrp2*$^{+/+}$ FVB/N: n = 4

(**G**) Immunofluorescence confocal microscopy detected PTTG1 in mitotic NIH-3T3 cells at the perinuclear region, concentrated at the centrosomes (**a**, arrowheads) positive for γ-tubulin and in the cytoplasm (**a**, asterisk). Quiescent/interphase cells (**b**, **c**) showed PTTG1 localized to primary cilia stained with ARL13b. PTTG1 signals were detected at the base (**b**), as well as along the ciliary shaft (**c**). Insets show the magnification of the chosen cilia (dashed line squares). Experiments were repeated at least five times in triplicates. Scale bars: 1 µm.

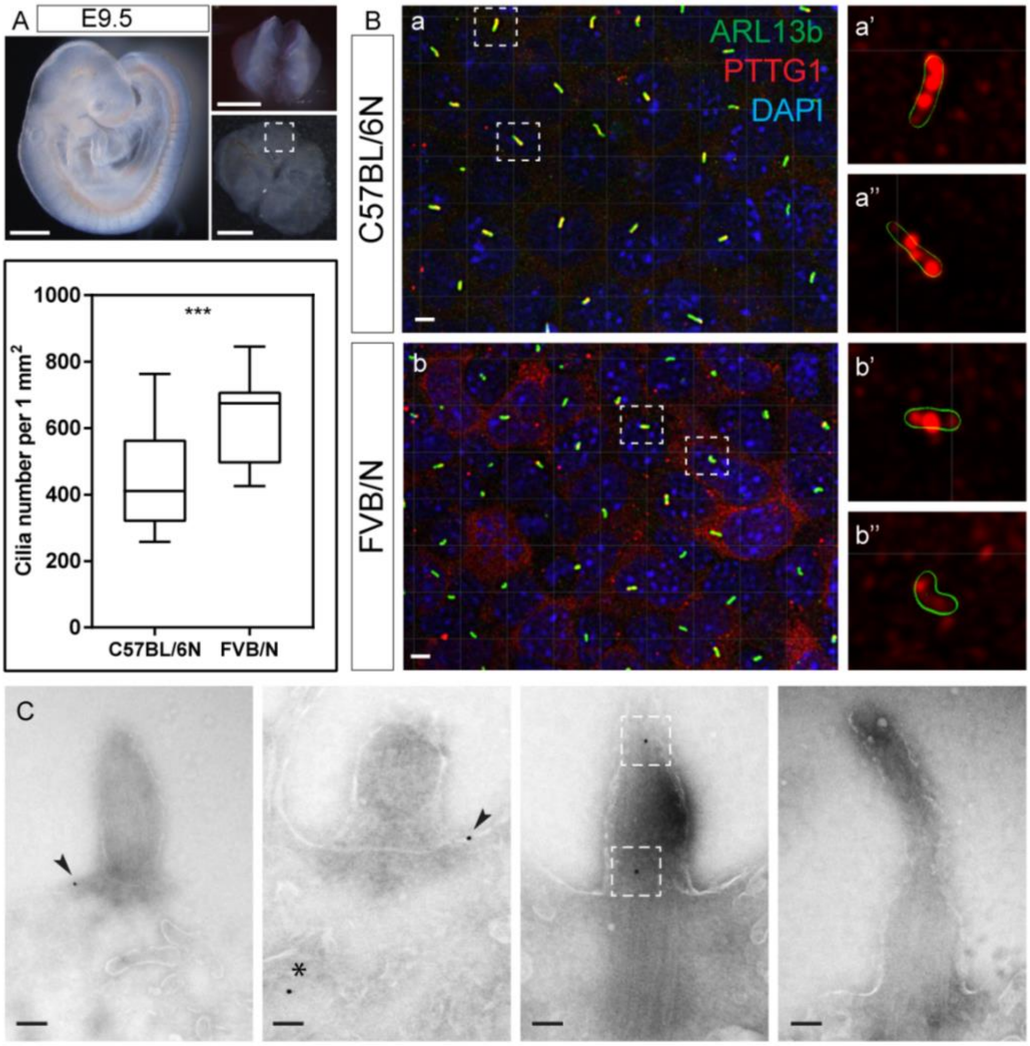


**Figure S6: Analysis of cilia and ciliary PTTG1 localization. Related to Figure 6**

(**A**) E9.5 embryos were used to prepare cephalic explants in order to examine the primary cilia of the mouse neuroepithelium. Images show the side view on the E9.5 embryo, head after opening the neural tube and flattened explants ready for the immunofluorescence staining. Dashed line square indicates the forebrain region of interest. Scale bars: 500 µm.

(**B**) Confocal microscopy images of the en face view on the neuroepithelium from *Lrp2*$^{+/+}$ C57BL/6N (**a**) and FVB/N (**b**) embryos showed two aspects: 1) PTTG1 localization to a subset of cilia in both strains; 2) differences in the cilia number per neural tube area between the two strains. Two representative primary cilia for each strain are shown in higher magnification (**a'**, **a''** and **b'**, **b''**) and displayed PTTG1 localization to the ciliary shaft for both backgrounds. Green reconstructed ARL13b staining showed the circumference of each cilium. Scale bars: 1 µm.

The graph represents the quantification of the average cilia number per 1 mm$^2$ area with whiskers indicating minimal and maximal values. N = 5 C57BL/6N and n = 6

FVB/N embryos were analyzed and a total number of n = 1636 and n = 1742 cilia, respectively, was quantified. Unpaired T-test statistical analysis was performed; *** p < 0.0001.

(**C**) Immunogold labeling of PTTG1 in the primary cilium of E9.5 C57BL/6N neuroepithelium. PTTG1 localized to a subset of primary cilia and was found in periciliary region (arrowheads), the ciliary shaft (boxed) and in the cytoplasm (asterisk), confirming the observations from the confocal microscopy. Representative images show cilia positive for PTTG1 (three left panels) and one cilium without PTTG1 signals (right panel). 2 embryos were analyzed for immunogold labeling; experiments were repeated 5 times with a number of 40 sections analyzed for each experiment. Scale bars: 100 nm.
